# A cellulose synthase interactome uncovers BAG proteins as regulators of cellulose synthase homeostasis

**DOI:** 10.64898/2026.09.28.755015

**Authors:** Manoj Kumar, Simon Turner

**Affiliations:** School of Biological Sciences, The University of Manchester, Manchester, UK

## Abstract

Cellulose synthase complexes build the load-bearing cellulose microfibrils of plant cell walls, yet how the abundance of their catalytic CELLULOSE SYNTHASE A (CESA) subunits is maintained remains unclear. Here, we used multi-bait TurboID proximity labelling with ten cellulose-synthesis-associated baits and six subcellular controls to define a high-confidence cellulose synthase neighbourhood. Stringent spatial and recurrence-based filtering yielded a core network of 119 interactions among 44 proteins and identified three members of the conserved Bcl-2-associated athanogene (BAG) family as previously unrecognised regulators of cellulose synthase homeostasis. BAG1-3 associated with primary-wall CESAs in reciprocal proximity-labelling experiments. Arabidopsis bag mutants showed reduced cellulose accumulation, hypersensitivity to cellulose-synthesis inhibitors, and markedly decreased CESA protein abundance without corresponding changes in CESA transcript levels. Loss of BAG function also increased the accumulation of CESA6 in vacuolar compartments. These findings identify BAG proteins as previously unrecognised regulators of cellulose synthase homeostasis and link a conserved proteostasis-associated protein family to plant cell wall biosynthesis. More broadly, the study establishes multi-bait proximity labelling, combined with cell location-specific controls, as a strategy for resolving dynamic protein networks whose components traffic through multiple subcellular compartments.

## Introduction

Cellulose microfibrils are the principal load-bearing polymers of plant cell walls and are synthesised at the plasma membrane by cellulose synthase complexes (CSCs). In land plants, CSCs are rosette-shaped assemblies built from CELLULOSE SYNTHASE A (CESA) catalytic subunits, whose spatial organisation is thought to influence glucan-chain assembly and, ultimately, cellulose microfibril structure and material properties ^1,2^. During primary wall synthesis, CSC isoforms are delivered to the plasma membrane, move along cortical microtubule trajectories and synthesise cellulose as cells expand. Thus, cellulose production depends not only on CESA catalytic activity but also on the cellular mechanisms that assemble, deliver, position, recycle and remove CSCs. Live-cell imaging and genetics have identified several factors that influence CSC behaviour. These include proteins implicated in CSC assembly, microtubule guidance, stress-dependent maintenance of cellulose synthesis, exocytosis, endocytosis and recycling, such as STELLO proteins, CSI/POM2, CC proteins, PATROL1, exocyst components, clathrin-mediated endocytic machinery and SHOU4/SHOU4L ^3–9^. Despite this success and the crucial importance of cellulose synthesis in determining growth ^10^, many questions remain regarding how the abundance and lifetime of CESA proteins are regulated across the plasma membrane, endomembrane trafficking and degradation pathways. Similarly, the accumulation of defective CSCs in the plasma membrane can decrease cellulose synthesis, but how defective complexes are identified and removed from the plasma membrane, and their subsequent fate, remain unknown ^11,12^.

Defining this regulatory neighbourhood is technically challenging. Yeast two-hybrid and related binary assays test direct interactions but do not preserve the membrane, cytoskeletal and trafficking context in which CSCs operate. Affinity purification coupled to mass spectrometry can identify stable complexes, but low-abundance, membrane-associated, insoluble, weak or transient interactions are often lost during extraction. Proximity labelling provides a complementary strategy because an engineered enzyme fused to a bait protein covalently labels nearby proteins in living cells, enabling stringent purification and mass spectrometry-based identification of local protein neighbourhoods ^13–16^. TurboID has extended proximity labelling to plant systems, enabling detection of low-abundance protein complexes and cell-type- or compartment-specific proteomes in vivo ^14,16,17^. Nevertheless, most proximity-labelling studies have relied on one or a few baits, making it difficult to distinguish bait-specific proximity from general subcellular background. A multi-bait strategy offers a more powerful way to interrogate a large, dynamic and spatially distributed complex: prey recurrently detected across multiple CSC-associated baits are more likely to reflect the CSC pathway, whereas compartment-specific controls help remove proteins labelled simply because they occupy the same organelle, membrane or cytoskeletal environment. This logic has been demonstrated in other systems, where multiplexed proximity biotinylation converted individual bait datasets into a network-level view of complex organisation ^18^.

Here, we apply TurboID proximity labelling to a panel of CSC-associated bait proteins and compartment-specific controls to define a high-confidence CSC proximity interactome. By integrating multiple bait perspectives with spatial controls, we identify proteins associated with CSC assembly, trafficking and plasma-membrane function, and prioritise candidates for functional analysis. We then use this resource to uncover previously unrecognised regulators of CESA protein abundance, thereby establishing multi-bait proximity labelling as a discovery framework for cellulose synthesis and CSC regulation.

## Results

### Identifying a comprehensive interactome involved in cellulose biosynthesis

To overcome the limitations of using a single bait protein, we fused the TurboID tag to 10 proteins known to play a role in cellulose synthesis in the primary cell wall (Table S1, Figure 1A). All fusion proteins were functional and significantly complemented their respective mutants (Figure S1). The cellulose synthase complex exhibits a complex localisation pattern involving the Golgi, plasma membrane and small vesicles (MASCs) and is associated with cortical microtubules (Figure 1A). Consequently, we included TurboID: YFP-tagged control lines for soluble proteins (YFP), plasma membrane (PIP2a), Golgi (BET12), ER (STIM1), microtubules (MAP4) and actin cytoskeleton (LifeAct) (Table S1). After 7 days of growth on agar plates, seedlings were treated with 0.5 mM biotin for 3 hours to biotinylate proximal proteins (Figure S2). Biotinylated proteins were purified on streptavidin beads and subjected to on-bead digestion before mass spectrometric (MS) analysis. The MS data comprised several thousand spectra with associated signal intensities and peptide sequences. Based on peptide-to-protein inference, the data were separated into exclusive peptide data and ambiguous peptide data (Supplementary Dataset 1). Throughout the manuscript, unless otherwise stated, we refer to the exclusive peptide data. The numbers of peptide and protein identifications followed a similar pattern (Figure S3). A PCA plot based on exclusive peptide data shows tight clustering of the replicates, demonstrating the high biological reproducibility of the mass spectrometry data (Figure S4). Furthermore, the distinct spatial separation between the CSC test baits and the control groups underscores the specificity of the proximity labelling approach and validates the distinct proteomic micro-environments captured by each bait category.

**Figure 1.**
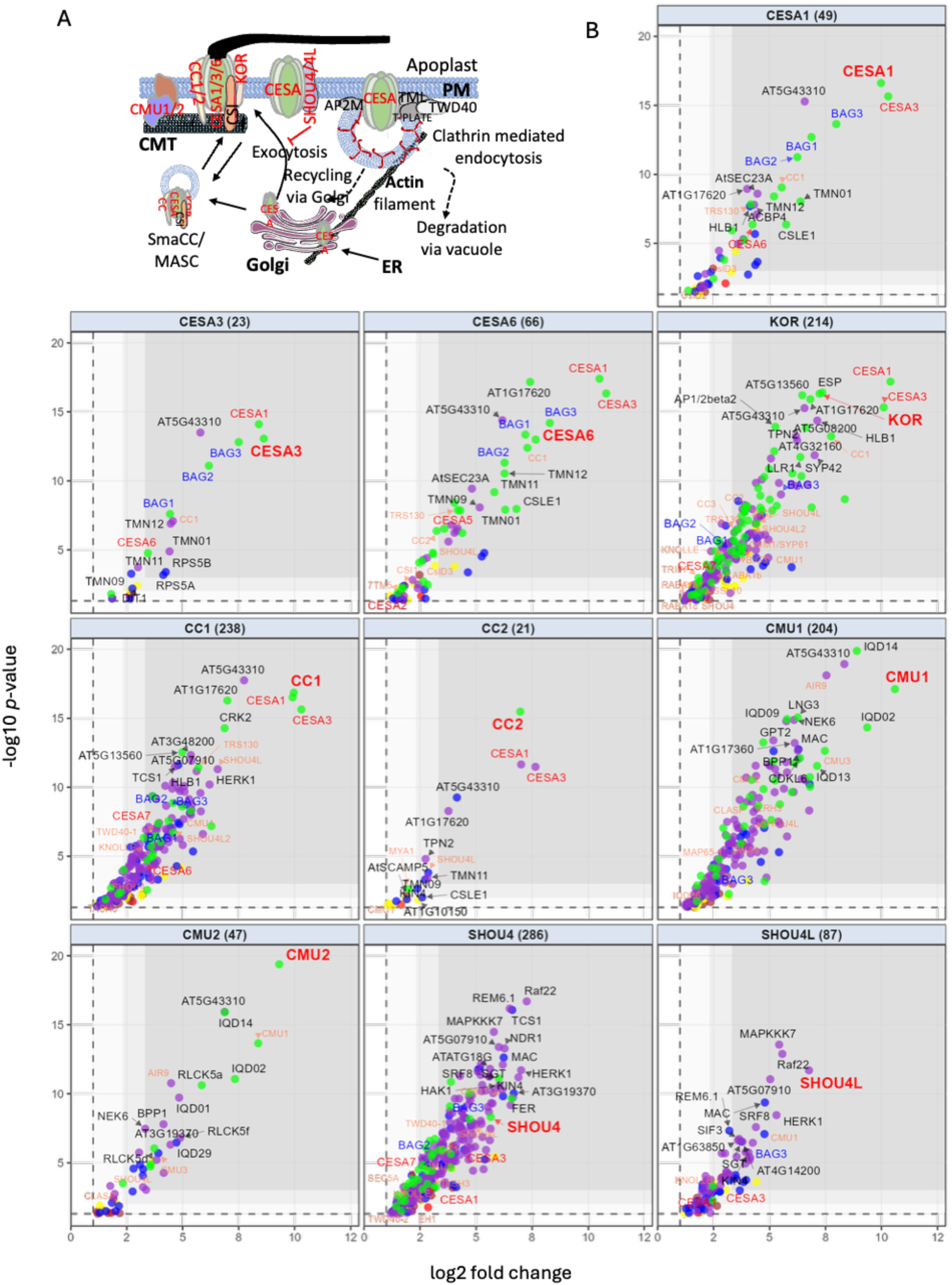
Significant interactions identified by all baits. **A.** Schematic model of Cellulose Synthase Complex (CSC) intracellular trafficking and selection of proximity labelling baits. Test baits are highlighted in red, control baits with bold text (ER – STIM1, Golgi – BET12, PM – PIP2a, CMT – MAP4 and Actin-lifeAct). Abbreviations: CSC, Cellulose Synthase Complex; PM, Plasma Membrane; ER, Endoplasmic Reticulum; CMT, Cortical Microtubules; SmaCC/MASC, Small CESA Compartments / Microtubule-Associated Cellulose Synthase Compartments. **B.** Multi-panel volcano plots illustrating the differential protein enrichment for bait proteins versus the EYFP control. The grid displays the enrichment profiles for the 9 of 10 test baits. The total number of interactors for each bait is provided in parentheses within the panel headers. For each panel, proteins are plotted by their log2 fold change on the x-axis and -log10 p-value on the y-axis. Data points are colour-coded based on the number of organelle marker baits (out of 5) against which they remained significantly enriched: 0 (red), 1 (brown), 2 (yellow), 3 (blue), 4 (purple), and 5 (green). The bait protein is highlighted in large red text; CESA proteins are in red; BAG family proteins are in blue; and known cellulose-synthesis-related proteins (as defined in Gu and Rasmussen, 2022) are highlighted in salmon.

A total of 5266 interactions involving 1822 proteins were detected across the 16 baits (10 test baits and 6 compartment-specific control baits) (Table S2). Complexity was significantly reduced by removing interactions not enriched relative to the soluble YFP control bait, leaving 2784 interactions among 1333 proteins (Table S2, Figure 1B and S5). Volcano plots for individual baits, based on this dataset, demonstrate that we can identify a large number of components that have at least some role in cellulose synthesis^8^ and capture known interactions between CMU1 and IQD proteins, which are not necessarily ascribed a role in cellulose synthesis ^19^ (Figure 1B). However, for proteins not previously associated with cellulose synthesis, it is difficult to distinguish specific interactions from those involving abundant proteins from the same compartment, which is particularly problematic for bait proteins known to reside in multiple cellular compartments.

### Identifying a core interactome

When we performed a similar pairwise comparison of compartment-specific control baits against YFP, we enriched for other known markers of those compartments (Figure S5). However, to better visualise the data, we generated a network from these 2784 interactions and colour-coded each protein according to its location in the SUBA database ^20^. In general, proteins of known location grouped with the control bait for that location (Figure S6). Enrichment of the bait proteins against the 5 remaining control baits further reduced the interactome to 227 interactions involving 152 proteins. The network appeared to separate into a core associated with the cellulose synthesis complex and a subnetwork comprising SHOU4 and CMU1, which play roles in inhibiting endocytosis and stabilising microtubules (Figure 2A). This was confirmed by hierarchical clustering of the dataset, which was consistent with known functions: all three CESA proteins, which together form a complex, clustered closely together but were clearly separated from SHOU4 and CMU1 (Figure 2B). The interactome revealed that some bait proteins produced numerous unique interactions identified by only a single bait (Figure 2C). For example, KOR, SHOU4 and CMU1 have 49, 29 and 26 interactions that were identified by no other baits (Figure 2C). In the case of KOR, it may have other functions within the cell apart from its role in cellulose synthesis; however, KOR also exhibits a unique trafficking pathway that involves trafficking via the vacuolar membrane ^21,22^. Since the focus was to identify a core network involved in cellulose synthesis, we exploited our multi-bait strategy to overcome this problem and included only proteins enriched against all controls and identified by 2 or more baits. This higher-confidence core network (Figure 3) retains a similar overall structure to the previous network (Figure 2A) but is reduced to 119 interactions between 44 proteins, composed of 10 bait proteins and 34 potentially novel interactors.

**Figure 2.**
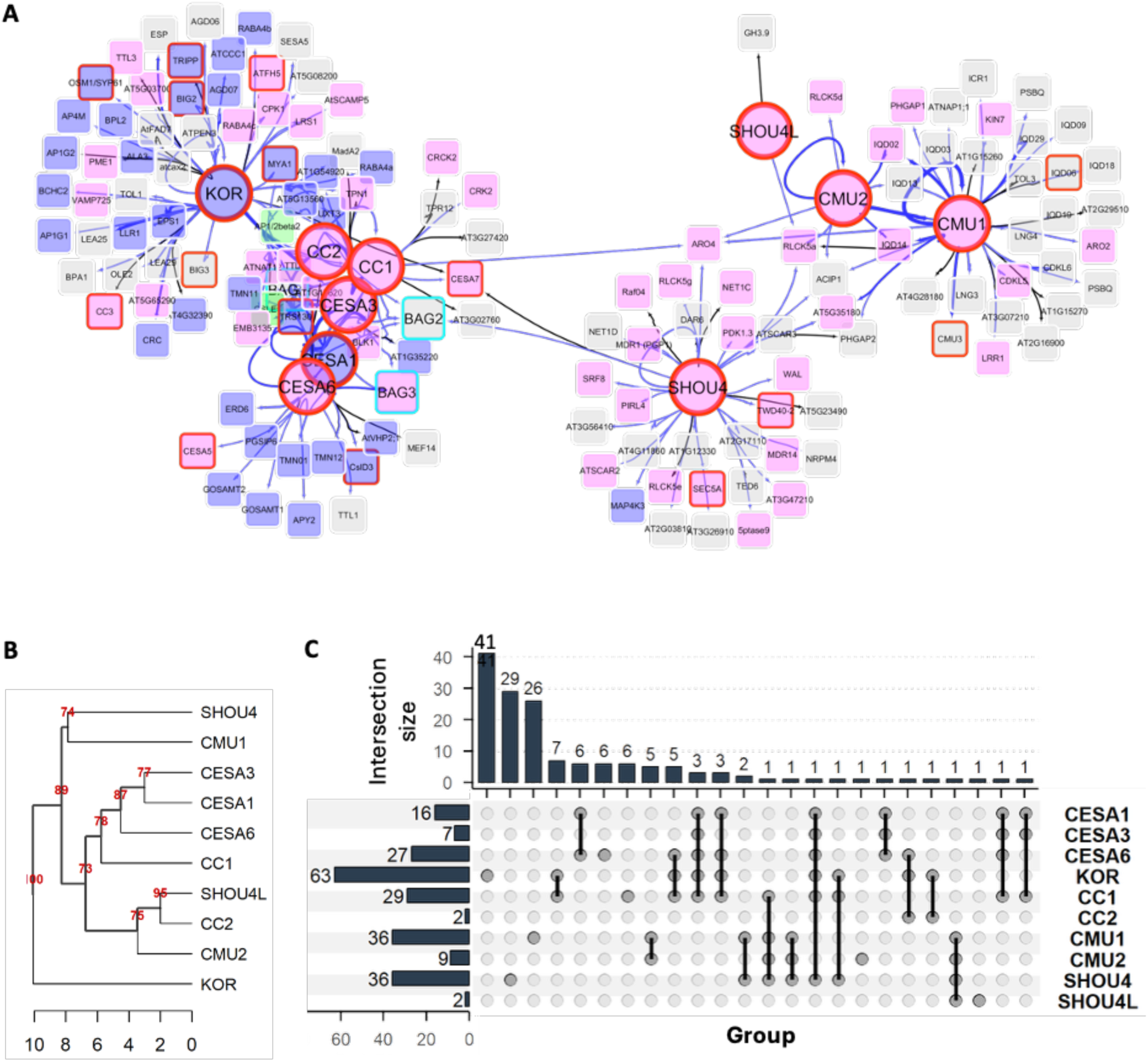
The core Cellulose Synthase Complex (CSC) interaction network. **(A)** A network graph visualising the core proximity interactions identified following stringent spatial filtering (significantly enriched against NoBait, EYFP, and all five organelle control baits). The central nodes with red borders denote the individual proximity labelling baits, which are connected via edges to their respective high-confidence interactors. **(B)** Hierarchical clustering dendrogram of the CSC baits based on the similarity of their proximity interaction profiles from the core network in (A). Red numbers at the nodes indicate bootstrap confidence values, demonstrating the robust functional and spatial grouping of the baits. **(C)** An UpSet plot detailing the overlap and uniqueness of the interactomes across the different baits. The horizontal bars (’Set size’) indicate the total number of core interactors identified for each individual bait. The vertical bars (’Intersection size’) quantify the number of shared or unique interactors corresponding to the specific combinations of baits indicated by the connected dots below the plot.

**Figure 3.**
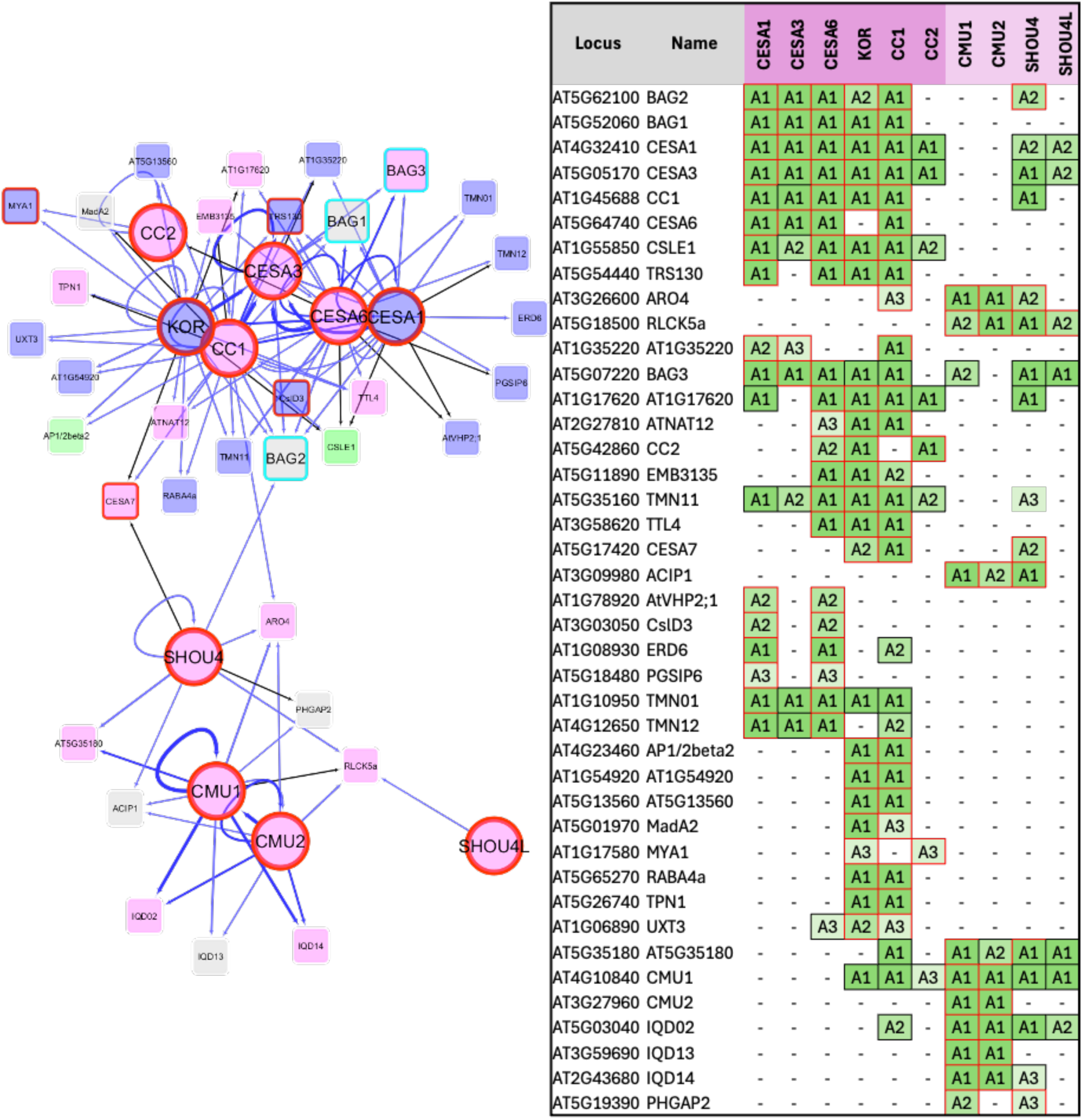
The highest-confidence consensus core interactome. **(A)** A network graph visualising the highest-confidence subset of the core CSC interactome, specifically highlighting robust, highly connected interactors identified by two or more independent proximity labelling baits. The central bait proteins are denoted by red borders. **(B)** A comprehensive interaction matrix detailing the specific bait-interactor relationships for all proteins displayed in the network in (A). The ten independent CSC baits are arrayed across the columns. The text within the green shaded boxes (A1, A2, A3) denotes the relative magnitude of statistical enrichment compared to the EYFP control (corresponding to high, medium, and low enrichment levels, respectively). The border colour of each cell defines the spatial filtering stringency of that specific interaction: red borders indicate enrichment against all five organelle control baits, whereas black borders indicate interactions enriched against three or four of the organelle controls.

The core network was enriched for proteins associated with the trans-Golgi network and the Golgi apparatus (Figure 3A and S7), including members of the transmembrane nine (TMN) family of Golgi-resident proteins ^23^ and the small GTPase RABA4a, which is linked to cell wall deposition ^24,25^. Among the proteins identified were CESA7, associated with cellulose synthesis in secondary cell walls, and CSLD3, which synthesises cellulose in root hairs ^26,27^. Although these proteins could form mixed complexes with the CESA proteins used here, shared plasma membrane localisation and trafficking pathways may be sufficient to account for their enrichment in our analysis. Furthermore, some proteins, such as KOR, play a role in cellulose synthesis in both primary and secondary cell walls^28^. Other bait proteins may also play a similar role, resulting in the interactome containing additional proteins involved in secondary cell wall biosynthesis.

### BAG1-3 localise to the plasma membrane but are not integral components of the cellulose synthase complex

To verify that our interactome was robust and capable of identifying novel proteins required for cellulose synthesis, we focused on members of the Bcl-2–associated athanogene (BAG) family. Mammalian BAG proteins play a pivotal role in determining cell fate, functioning as co-chaperonins with Hsp70 to regulate proteostasis ^29^. The Arabidopsis genome encodes 7 BAG proteins, which are divided into 2 groups (Figure S8A). BAG1-4 all possess a BAG domain and a ubiquitin-like domain, a structure that closely resembles their mammalian homologues. BAG4-7 lack the ubiquitin-like domain and are much more variable in length (Figure S8A). However, they have been shown to be important in resistance to a variety of biotic and abiotic stresses ^30^. In this high-confidence network, BAG1, BAG2 and BAG3 were enriched by 3-6 test baits, including all 3 CESA proteins, suggesting a strong association of BAG1, BAG2 and BAG3 with the cellulose synthase complex (Figure 3).

We used live-cell imaging to examine the relationship between BAG and CESA proteins in more detail, focusing on stably transformed YFP-tagged BAG1 in Arabidopsis seedlings. BAG3 has been reported to localise to the cytoplasm ^31^. However, our data suggest that BAG1 exhibits a punctate pattern at the cell surface (Figure 4A), similar to that recently reported for BAG2. This pattern is similar to that described for CESA proteins ^32^. However, we saw no evidence that BAG1 formed tracks in the plasma membrane, and kymograph analysis of live-cell imaging suggested that BAG1 was stationary at the plasma membrane (Figure 4B). This contrasts with CESA6, which follows tracks in the plasma membrane as the cellulose is consistent with several studies using immunoprecipitation with CESA that have failed to identify any BAG proteins ^33,34^ and suggests the interaction is likely to be weak or transient.

**Figure 4.**
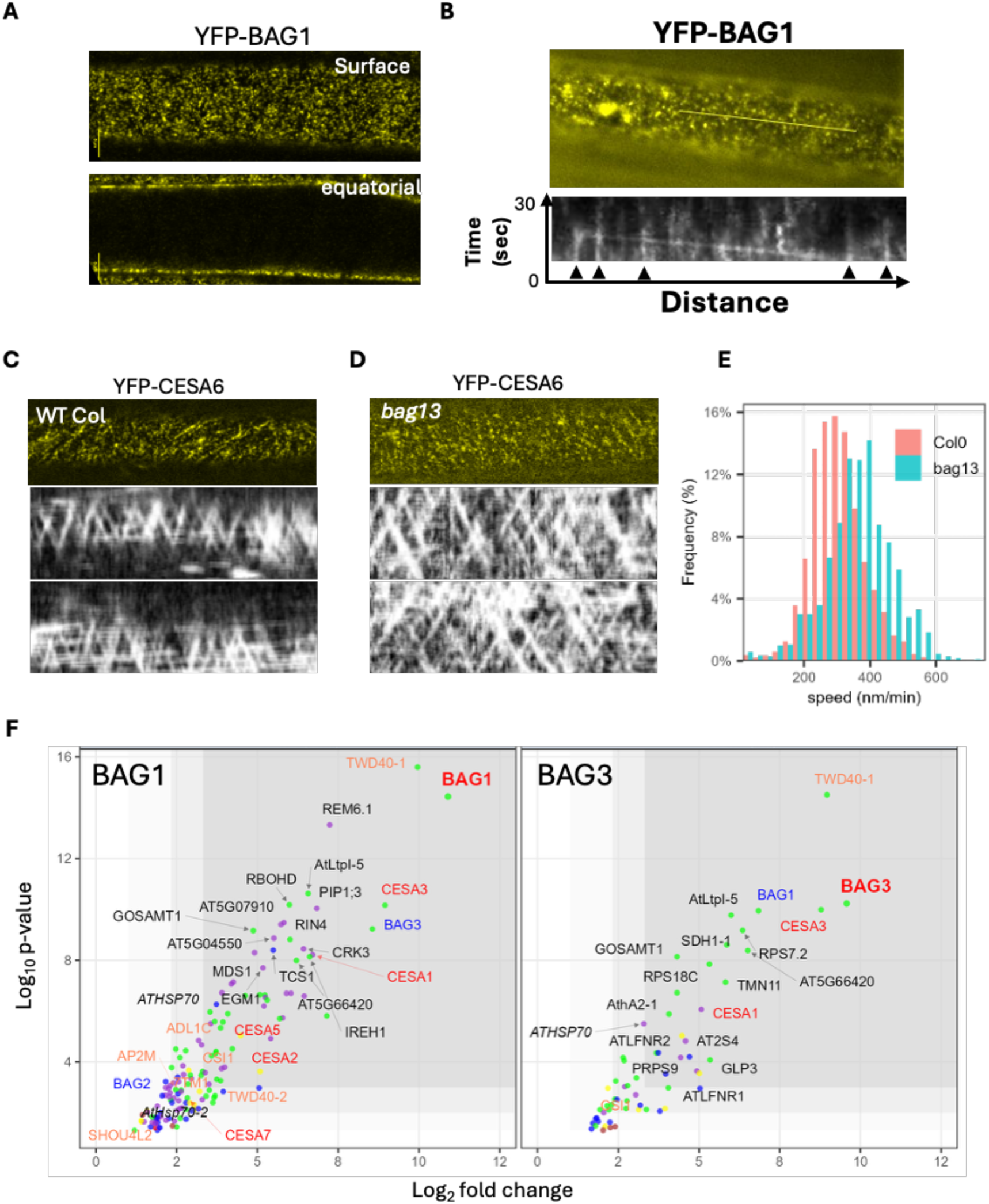
BAG proteins associate with CESA proteins but are not part of the cellulose synthase complex. **(A-B)** Subcellular localisation of YFP-BAG1 in root epidermal cells, using confocal microscopy (A) or etiolated hypocotyls using TIRF microscopy (B). **(C-E)** Analysis of Cellulose Synthase Complex (CSC) dynamics. **(C, D)** Representative confocal images at the plasma membrane focal plane of epidermal cells of etiolated seedlings expressing YFP-CESA6 together with representative kymographs generated from time-lapse imaging in either wild type (Col0) **(C)** or bag1,bag3 double mutants **(D)**. **(E)** Frequency distribution histogram quantifying the migration speed of YFP-CESA6 particles (nm/min in the bag13 mutant (blue) and Col0 (red). **(F)** Volcano plots illustrating the dìerential enrichment analysis for the BAG1 and BAG3 test baits compared to the EYFP soluble control. Data points are colour-coded based on the number of organelle marker baits (out of 5) against which they remained significantly enriched: 0 (red), 1 (brown), 2 (yellow), 3 (blue), 4 (purple), and 5 (green). The bait protein is highlighted in large red text; CESA proteins are in red; BAG family proteins are in blue.

As we were unable to confirm the CESA-BAG interaction by immunoprecipitation, we generated TurboID-tagged versions of BAG1 and BAG3 to obtain independent evidence of in vivo proximity and verified their functionality by complementing the *bag1,3* double mutant (Figure S11). We then performed proximity labelling and mass spectrometry using the same protocol as for other baits in this study. The volcano plots for BAG1 and BAG3 (Figure 4F) show that, consistent with its plasma membrane location, BAG1 is enriched for several known plasma membrane markers, including a remorin, PIP1 and RIN4. However, none of these proteins was enriched relative to our PIP2 control bait. In contrast, when using BAG1 as bait, CESA3 and BAG3 were among the top 3 most enriched proteins and were enriched relative to all control baits. Similarly, CESA3 and BAG1 were among the three most enriched proteins when using BAG3 as bait and were significantly enriched against all controls. Other primary cell wall CESAs were also enriched using BAGs as baits, confirming that, at some location within the cell, the CSC and specifically CESA proteins exhibit a close association with BAG proteins in vivo (Figure 4F).

### Plant BAG proteins as regulators of CESA protein abundance

The aim of this study was to identify novel components required for cellulose synthesis. To determine whether any of the novel proteins within our network had important functions in cellulose synthesis, we identified T-DNA mutants likely to be knockouts in 20 of the 33 genes previously not assigned a role in cellulose synthesis, and tested the effects of the cellulose synthase inhibitor isoxaben. We found that single mutants exhibited only marginal effects on hypocotyl elongation, and none were comparable to the moderate cellulose-deficient allele *rsw1-1* (Figure S12). However, given that some families, such as the BAG proteins, contained several closely related proteins that formed part of the network, we examined whether functional redundancy was masking a clearer phenotype. Consequently, we generated all double-mutant combinations and the *bag1*, *bag2*, *bag3 triple mutant*. Of the single mutants, only *bag2* was 20% shorter than the wild type, while *bag1* and *bag3* exhibited no phenotype. In contrast, the *bag1, bag3* double mutant and the *bag1, bag2, bag3* triple mutant were significantly shorter than the wild type (Figure 5A). The phenotype was further enhanced when seedlings were grown in the presence of the cellulose synthase inhibitors isoxaben or DCB, with the *bag1, bag3* double and the *bag1, bag2, bag3* triple mutants reaching only a quarter of the length of the wild type, comparable to the phenotype of the known cellulose-deficient *rsw1-1* allele caused by a mutation in CESA1 ^35^ (Figure 5A). The bag triple mutants did not exhibit any decrease in cell number in the hypocotyl length (Figure S13) but showed shorter, swollen cells in the presence of DCB. This increased sensitivity to cellulose synthase inhibitors in the *bag1, bag3* double and *bag1, bag2, bag3* triple mutants correlated with a significant reduction in cellulose content in 7-day-old seedlings (Figure 5B). The reduction of around a quarter exhibited by the triple mutant is comparable with previously described CESA protein mutant alleles, *rsw1-1* ^35^ (Figure 5B). We also observed a comparable reduction in cellulose content in *bag1, bag3* and *bag1, bag2, bag3* multiple mutants in developing and mature inflorescence stems (Figure 5C,D).

**Figure 5.**
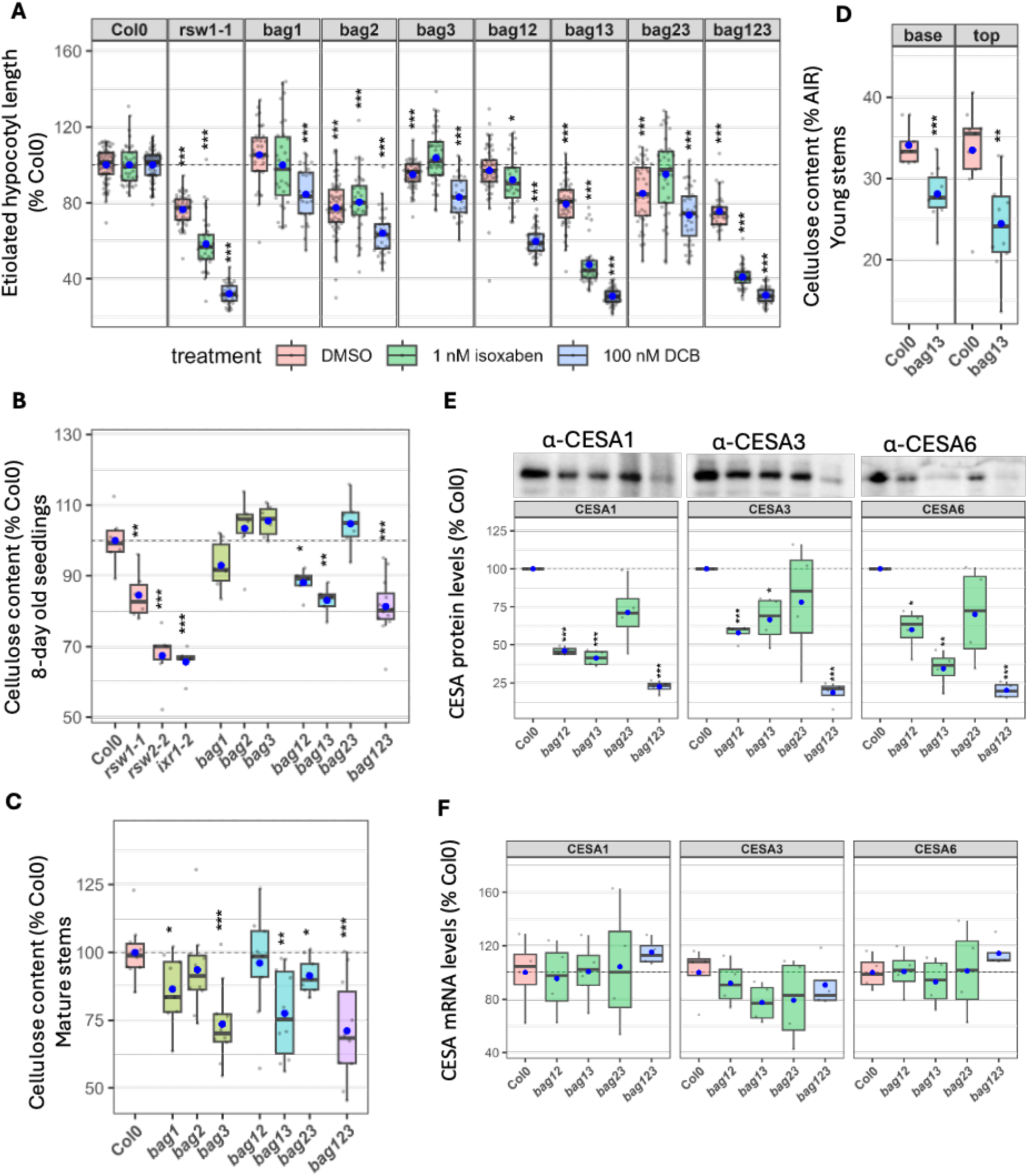
Bag mutants exhibit cellulose deficient phenotypes. **(A)** Quantification of etiolated hypocotyl length. Wild-type (Col-0), bag single, double, and bag1/2/3 triple mutants were grown together with known cellulose deficient mutants (rsw1, rsw2, ixr) and wild-type seedlings on isoxaben or 2,6-dichlorobenzonitrile (DCB). **(B–D)** Crystalline cellulose content of 8-day-old seedlings **(B)**, mature stems **(C)** and developing stems from 5-week-old plants **(D). (E)** Western blot analysis of primary wall CESA protein levels. Representative immunoblots probed with isoform-specific antibodies (α-CESA1, α-CESA3, and α-CESA6) are shown (top). **(F)** Quantitative RT-PCR (qPCR) analysis of CESA1, CESA3, and CESA6 transcripts. For all boxplots, the horizontal line represents the median, the box spans the interquartile range (IQR), and grey dots represent independent biological replicates. Asterisks indicate statistically significant differences compared to Col0. * p<0.05, ** p < 0.01, *** p < 0.001.

The well-established role of BAG proteins as co-chaperonins in mammalian and yeast cells ^36,37^ prompted us to test whether bag mutants affect the stability of CESA proteins. *bag1, bag3* and *bag1, bag2* double mutants, and the *bag1, bag2, bag3* triple mutants, all exhibited significantly reduced levels of CESA1, CESA3 and CESA6, with the *bag1, bag2, bag3* triple mutant possessing only 20% of the levels of all three CESA proteins compared to the wild type (Figure 5E). In contrast to the drastically reduced protein levels, CESA mRNA abundance remained stable and largely unchanged across all bag mutant backgrounds (Figure 5F), indicating that the effects on CESA protein levels were post-transcriptional and identifying BAG proteins as important regulators of CESA protein abundance.

### BAG mutants regulate vacuolar degradation of CESA proteins

Under normal growth conditions, YFP-tagged proteins cannot be tracked through the vacuolar degradation pathways because the low vacuolar pH quenches fluorescence. Fluorescent proteins can be visualised within the vacuole either by treatment with Concanamycin A to prevent proper localisation of the vacuolar ATPase, or by dark treatment, both of which increase the vacuolar pH ^38^. We focused on dark treatment to quantify the effects. We imaged YFP-CESA6 in WT and *bag1,bag3* double mutant at time zero, then after 3 and 6 hours of dark treatment (Figure 6A, B and S14). At time zero, fluorescence was apparent only at the plasma membrane or Golgi. However, by 3 hours we observed intracellular accumulation of YFP-CESA6 signal in structures that appeared to be pre-vacuolar compartments or lytic vacuoles, identical to those observed during similar experiments designed to look at the degradation of the boron transporter (BOR1) or the auxin transporter PIN1 in the presence of boron or auxin, respectively (Figure 6A, S14). ^38,39^These intracellular compartments were more numerous at 6 hours and were more apparent in the bag double mutant, even though the overall signal in the double mutant was much lower (Figure 6A,B, S14). To quantify these results, we performed segmentation analysis of these structures (Figure 6C, S15). We observed an increase in the number of these vacuoles in the *bag1,bag3* double mutant following both 3 and 6 hours of dark treatment (Figure 6C). ConcA treatment gave a similar result with YFP fluorescence, with intracellular compartments being more apparent in the *bag1,bag3* double mutant than in the wild type (Figure S16).

**Figure 6.**
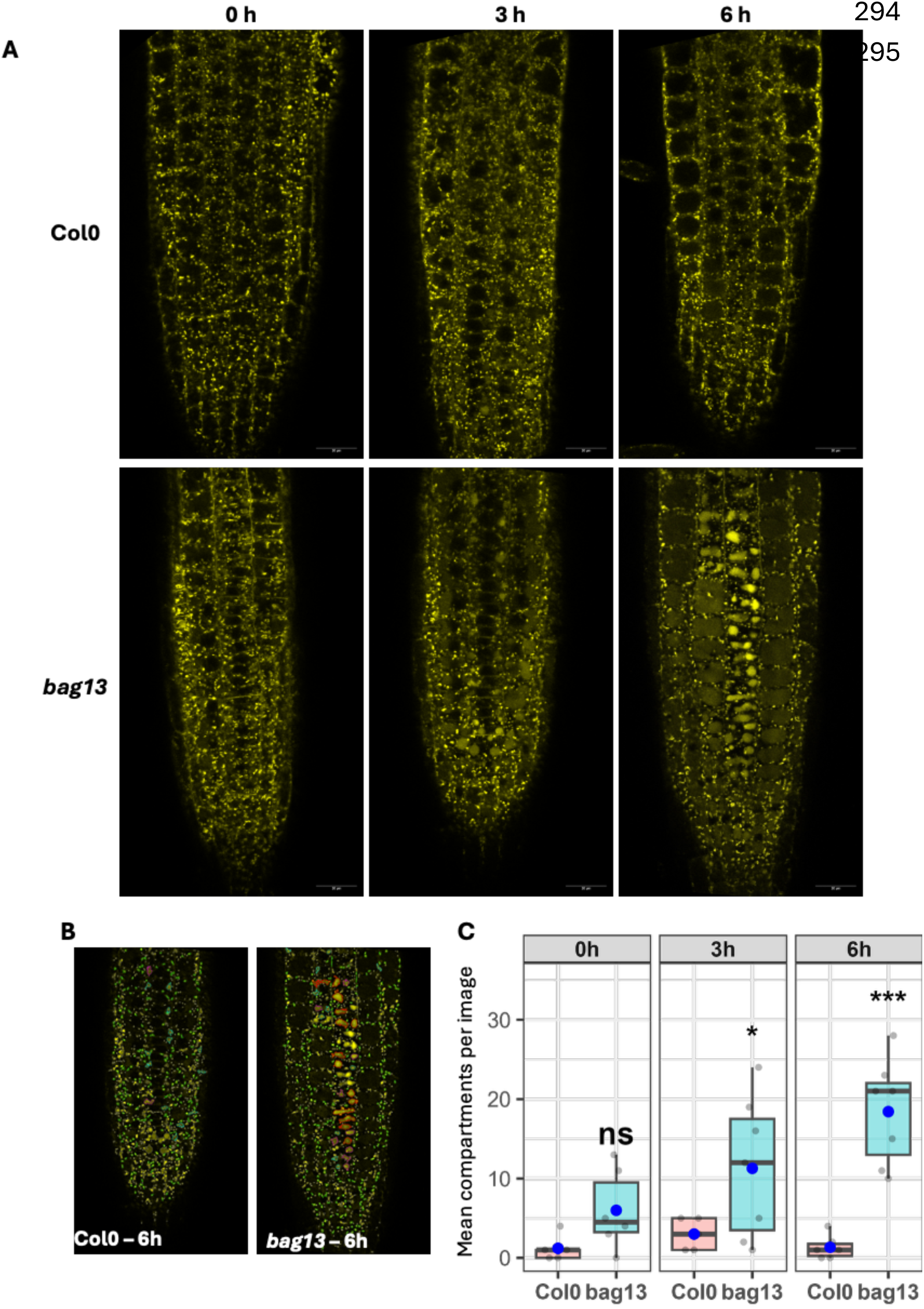
Quantitative analysis of YFP-CESA6 in dark-treated Col0 and bag13 mutant. **(A,B)** Examples of images from 5-day-old wild-type (Col0) and bag13 double mutant seedlings after 0, 3 and 6 hours of dark treatment **(A)**, alongside computational segmentation masks for 6h images **(B)**. The masks demarcate distinct intracellular structures used for automated particle classification. **(C)** Quantification of total macroscopic YFP-CESA6 defined intracellular compartments per root tip across the 0, 3, and 6-hour dark treatment time course. The horizontal black line represents the median, the blue dot indicates the arithmetic mean, and translucent grey points represent individual biological measurements. Asterisks denote statistical significance (* p < 0.05, p < 0.01, *** p < 0.001; ns = not significant).

## Discussion

Proximity-labelling approaches are increasingly used to map protein association networks in plants, yet their interpretation is often complicated by the large number of proteins that meet statistical enrichment criteria ^13,15^. This challenge is particularly acute for trafficking systems, where proteins occupy multiple intracellular compartments and suitable controls may not always be available. Here, we demonstrate that combining multiple cellulose-synthesis-associated bait proteins with compartment-specific controls is an effective strategy for distinguishing functional proximities from compartment-specific background labelling. By further requiring candidate proteins to be recovered by multiple independent baits, we reduced the number of detected proximities from more than 15,000 to a high-confidence network of 44 proteins. Importantly, this approach retained known components of cellulose biosynthesis ^1,8^ while identifying previously unrecognised regulators of cellulose synthase abundance (Figure 3).

A key advantage of the multi-bait strategy is that independent bait proteins provide orthogonal evidence for association with a biological process. Unlike networks derived from a single bait, inclusion in the final interactome depended not only on statistical enrichment but also on reproducible recovery across multiple components of the cellulose synthesis machinery. This approach helps overcome limitations arising from imperfect compartment controls and reduces the likelihood that proteins are retained simply because they are abundant components of a shared subcellular environment. Similar multi-bait strategies have proven powerful in mammalian systems ^18,40^, and our results demonstrate their utility for resolving complex membrane-trafficking networks in plants.

The identification of BAG proteins provides strong validation of the predictive power of this approach. BAG1–3 were recovered by multiple independent bait proteins, including all three primary cell wall CESAs, and showed some of the strongest enrichments in the dataset. Functional analysis demonstrated that BAG proteins are required for normal cellulose synthesis, as loss-of-function mutants exhibited reduced cellulose accumulation and substantially decreased levels of CESA proteins, despite largely unchanged transcript abundance. Together, these findings establish BAG proteins as important determinants of cellulose synthase abundance and reveal a previously unrecognised connection between BAG family proteins and plant cell wall biosynthesis. While BAG proteins are well established as co-chaperones involved in proteostasis in other systems ^36,41^, their role in cellulose biosynthesis has not previously been reported. An intriguing aspect of the BAG–CESA association is that it was readily detected by proximity labelling but not by immunoprecipitation. Previous studies investigating CESA-associated proteins using immunoprecipitation have similarly failed to identify BAG proteins ^33,34^. This discrepancy suggests that fBAG proteins may interact with cellulose synthase complexes only transiently or within spatially restricted compartments. Such interactions are inherently difficult to capture with conventional biochemical approaches and highlight one of the principal advantages of proximity-labelling technologies for studying dynamic trafficking pathways ^13^.

Our data support a model in which BAG proteins influence the post-transcriptional fate of cellulose synthase complexes. Loss of BAG function results in reduced CESA abundance and increased accumulation of CESA within vacuolar compartments, consistent with enhanced targeting of internalised complexes for degradation (Figure 6). However, the molecular basis of this regulation remains unclear. Proximity labelling using either BAG1 or BAG3 as bait identified a strong association with TWD40-1 and, to a lesser extent, with TWD40-2 (Figure 4F). Both proteins are components of the T-PLATE endocytic machinery, which has been implicated in cellulose synthase trafficking ^11,42,43^. These observations raise the possibility that BAG proteins influence trafficking decisions following CSC internalisation. Whether BAG proteins act directly in endocytic sorting, function through chaperone-mediated quality control pathways, or regulate CESA fate through an alternative mechanism remains an important question for future study.

In summary, this study demonstrates the power of multi-bait proximity labelling to define high-confidence interaction networks associated with complex membrane-trafficking pathways. Applying this approach to cellulose biosynthesis identified a robust core interactome and uncovered BAG proteins as previously unknown regulators of cellulose synthase abundance. These findings provide both a valuable resource for the community and new insight into the mechanisms that maintain cellulose synthase homeostasis in plant cells.

## Data Availability

The mass spectrometry proteomics data have been deposited to the ProteomeXchange Consortium via the PRIDE partner repository with the dataset identifier PXD080607. Reviewer access details: Log in to the PRIDE website using the following details: Project accession: PXD080607, Token: uwD1q9Lnz8Yl

## Methods

### DNA Cloning and plant transformation

Expression cassettes with a general structure, Promoter::Coding_Sequence:terminator, were assembled on a pCambia2300 backbone using restriction ligation, Gateway cloning (ThermoFisher Scientific, UK), and overlap expression PCRs ^44–46^. Gateway entry clones for KOR, CC1, CC2, CMU1, CMU2, SHOU4, and SHOU4L were described previously ^47^. Entry clones for CESA3 and CESA6 in pDONOR207 were a kind gift from the Staffan Persson laboratory. Entry clones for CESA1, TurboID, TurboID-EYFP, EYFP, STIM1_ER_reporter, BET12, PIP2A, MAP4_MBD, LifeAct, BAG1, BAG2, and BAG3 in the entry vector pDONOR/pZEO were generated in this study using primers described in Table S3. TurboID and miniTurbo sequences were codon-optimised for Arabidopsis, synthesised and cloned into pDONOR/pZEO. Coding sequences for Arabidopsis genes were amplified from Arabidopsis cDNA and cloned into pDONOR/pZEO. A MAPPER construct (STIM1_Signal_peptide-EGFP-STIM1_TMH-Linker_sequence-Polybasic_domain) was a generous gift from Yoshi Watanabe. The polybasic domain of this construct was replaced with TurboID by overlap extension, and the resulting construct was cloned into the entry vector pDONOR/pZEO.

Gateway destination vectors on the pCambia2300 backbone with native promoters pKOR, pCC1, pCC2, pCMU1, and pCMU2 were described previously ^47^. In this study, we constructed new destination vectors with native promoters pCESA1, pCESA3, pCESA6, pSHOU4, pBAG1, pBAG2, pBAG3, and pUBQ10. Some of these destination vectors included a TurboID(.EYFP) tag on the backbone, while others were untagged. The component fragments of these vectors (promoters, tags) were PCR-amplified and cloned into a pJET vector using a CloneJET PCR Cloning Kit (ThermoFisher Scientific, UK). All primers (Table S3) included appropriate restriction sites to allow concatenation of the fragments. The inserts in the pJET vector were fully sequenced before assembly to create the final destination vectors.

TurboID fusions were produced either by overlap extension PCR followed by LR reactions into untagged destination vectors, or by direct LR reactions into tagged destination vectors. Junction primers used to fuse genes and tags are listed in Table S3. The coding sequences were then verified by Sanger sequencing.

### Plant Material

Verified expression clones were transformed into Arabidopsis via Agrobacterium-mediated plant transformation using the floral dip method (Clough and Bent, 1998). The loss-of-function mutants used to transform test bait constructs, cesa1any1 (Fujita et al., 2013), ces3ixr1-2 (Scheible et al., 2001), cesa6prc1-1 (Fagard et al., 2000), kor1-1 (Nicol et al., 1998), cc1, cc2, cc1 cc2 (Endler et al., 2015), cmu1, cmu2, cmu1 cmu2 (Liu et al., 2016), shou4, shou4l, shou4 shou4l (Polko et al., 2018) have all been described previously. T-DNA mutants for bag1-1 (SM_3.15968), bag2-1 (GK-493F03) and bag3-1 (SALK_124153) and other candidates shown in Figure S12, at5g52060 (bag1) - SM_3_15968; at5g62100 (bag2) - GABI_493F03; at5g07220 (bag3) - SALK_124153; at3g26600 (aro4) - SALK_122722; at5g18500 - GABI_742B08; at1g17620 - SALK_003242; at1g10950 (tmn01) - GABI_284G01; at5g35160 (tmn11) - GABI_053H04; at4g12650 (tmn12) - GABI_760D08; at3g09980 (acip1) - SALK_028810; at3g58620 (ttl4) - SALK_033390; at1g17580 (mya1) - SALK_022140; at5g13560 - SALK_120883; at2g27810 (atnat12) - SAIL_764_H08; at3g03050 (csld3) - SALK_112105; at1g55850 (csle1) - SALK_011984; at1g06890 (uxt3) - SAIL_213_A09; at1g35220 - SALK_020945; at5g35180 - SAIL_829_G02; at5g65270 (raba4a) - SALK_046239 were obtained from NASC, and homozygous plants were identified by PCR-based genotyping using primers described in Table S3. Double and triple mutant combinations were generated by crossing single mutants and identifying double and triple homozygous plants.

### Plant growth and analysis

Arabidopsis plants were grown on plates and in soil under standard laboratory conditions, as described previously ^44,47,48^. Briefly, Arabidopsis seeds were surface-sterilised with 10% sodium hypochlorite, stratified for 2 days at 4°C, and plated on ½ MS plates with or without antibiotics. After 7 days in an incubator, seedlings were transplanted into 9 cm square pots containing a 1:1:5 mixture of perlite, vermiculite and compost. Plants were then grown for a further 6 weeks in soil under long-day conditions (16h/8h day/night, 22°C/18°C temperature and 80% humidity). For etiolated hypocotyl elongation assays, plates were incubated in light for 20 hours before being transferred to the dark and grown for 4-5 days.

For root and etiolated hypocotyl length measurements, plates were scanned on a flatbed scanner at 600 dpi. Roots and etiolated hypocotyls were then identified by manually drawing segmented lines along individual seedlings in ImageJ (Fiji). The segmented line regions of interest (ROIs) were measured using built-in functions. Data were expressed as % WT.

For cellulose content analysis of stems, 50 mm pieces from the primary inflorescence stem, starting 5 mm above the base, were harvested and stored in 70% ethanol. Cellulose content in these pieces was then measured using a variation of the Updegraff method ^49,50^. For seedlings, 7-day-old seedlings were frozen in liquid nitrogen and ground into a fine powder using a bead mixer-mill (Qiagen, Tissue Lyser II). Alcohol-insoluble residue was then prepared, and cellulose content was measured as above, except that centrifugation was used at each step to remove liquids rather than aspiration.

Data organisation and preliminary processing were performed in Microsoft Excel. Advanced statistical analysis and data visualisation were conducted using custom R scripts in RStudio, utilising the *ggplot2* package for graphics and *DescTools* for statistical modelling. Pairwise comparisons were assessed using Student’s t-tests. Multiple comparisons against a designated baseline control were analysed using a one-way Analysis of Variance (ANOVA) followed by Dunnett’s post hoc test, with significance set at p < 0.05.

### Affinity purification of biotinylated proteins

To screen lines for biotinylation levels, up to 10 independent lines per genotype were grown on plates as described above. After 7-8 days of growth, 10 mL of water containing 0.5 mM biotin was pipetted directly over the roots of each plate. Seedlings were then incubated under normal growing conditions for 20 hours. Plates were kept flat, rather than standing vertically, to prevent the biotin solution from running off. Seedlings were rinsed with water, dabbed dry with paper towels, and frozen for protein extraction and analysis. Following analysis of crude extracts by Western blotting, 3-4 independent lines were selected for further analysis based on biotinylation and/or complementation levels. For proximity labelling, T2 seed from selected lines was pooled and grown on plates as above. Three biological replicates were grown for each genotype, and biotin application was performed as described above, but the biotin incubation was limited to 3 hours.

To remove excess biotin from the samples, they were subjected to trichloroacetic acid (TCA)/acetone and chloroform/methanol precipitations as described ^47^. Frozen Arabidopsis tissue powder (∼400 mg) was suspended in 10 mL of ice-cold TCA/acetone solution (10% w/v TC, 10 mM TCEP in acetone) and incubated at −20 °C for 1 hour. Samples were centrifuged at 5,000 rpm for 10 min, and the supernatant was discarded. Pellets were washed twice with ice-cold acetone, centrifuged, and air-dried for 10 min. The resulting white pellet was resuspended in 5 mL of LB2 buffer (6 M urea, 2.5% SDS, 100 mM Tris-HCl pH 7.2, 150 mM NaCl, 5 mM EDTA, 1 mM PMSF) and incubated at room temperature for 20 min with rotation. Crude extracts were clarified by centrifugation and filtration through Miracloth.

The samples were then subjected to chloroform–methanol precipitation (CMP) by mixing with methanol (3 volumes), chloroform (1 volume), and water (4 volumes). After centrifugation, the upper phase was discarded, 4 volumes of methanol were added, and the mixture was incubated at −20 °C overnight. Following a further centrifugation, the pellets were washed with 80% methanol, dried, and resuspended in 2 mL of LB2 buffer. A 100 µL aliquot of the extract was set aside as an input control for Western blot analysis. Dynabeads C1 Streptavidin magnetic beads (75 µL beads per mL of starting tissue) were washed three times with LB1 buffer (100 mM Tris-HCl pH 7.2, 150 mM NaCl, 5 mM EDTA). CMP extracts were incubated with the beads overnight at room temperature with rotation. The beads were then washed three times with wash buffer (100 mM Tris-HCl pH 7.2, 500 mM NaCl, 0.1% SDS). After the final wash, 150 µL of the sample was collected for Western blot analysis. Biotinylated proteins from this sample were eluted by boiling in 1.5× Laemmli buffer containing 50 mM DTT and 2 mM biotin for 10 minutes. The eluate was analysed by Western blotting alongside the flow-through fraction and input controls to assess the efficiency of the affinity purification (Figure S2).

The remaining beads were washed three times with 50 mM ammonium bicarbonate (Ambic) and digested overnight at 37 °C with 0.8 µg of Trypsin/Lys-C mix (Promega V5071) in 200 µL of Ambic. The following morning, the flow-through fraction, containing peptides from biotinylated proteins, was collected. Peptides were reduced by adding 50 mM DTT and incubating at room temperature for 30 minutes. They were then alkylated with 100 mM iodoacetamide (final concentration) for 1 h at room temperature in the dark, followed by quenching with an additional 50 mM DTT. Samples were stored at −20 °C until peptide purification.

### Mass spectrometry and proteomics data analysis

Reduced and alkylated peptides were acidified by adding an equal volume of 0.1% formic acid. Acidified samples were loaded onto OLIGO R3 reverse-phase resin (Thermo Scientific, catalogue number 1133903), washed twice with 0.1% formic acid, and eluted with 200 µl of 30% acetonitrile + 0.1% formic acid. Peptides were then transferred to mass spectrometry vials and dried completely using a speed vac. Digested samples were analysed by LC-MS/MS using an UltiMate® 3000 Rapid Separation LC (RSLC, Dionex Corporation, Sunnyvale, CA) coupled to an Exploris 480 (Thermo Fisher Scientific, Waltham, MA) mass spectrometer. Peptide mixtures were separated on a 75 mm x 250 μm i.d. 1.7 µM CSH C18 analytical column (Waters) using a multistep gradient from 95% A (0.1% FA in water) and 5% B (0.1% FA in acetonitrile) to 7% B at 1 min, 18% B at 35 min, 27% B at 43 min, and 60% B at 44 min at 300 nL min-1. Peptides were selected for fragmentation automatically by data-dependent analysis.

Database searching and PSM (peptide spectrum match) processing were performed in Proteome Discoverer 2.2 (Thermo Fisher Scientific, UK). Two complementary search engines, Sequest™ HT (Thermo Fisher Scientific, UK) and Mascot (Matrix Science, UK), were used to search the data. Both engines were configured to search the Arabidopsis thaliana Araport11 protein database (48359 entries) with trypsin as the digestion enzyme. Sequest was set to a Precursor Mass Tolerance of 10 ppm and a Fragment Mass Tolerance of 0.02 Da, while Mascot used a Precursor Mass Tolerance of 20 ppm and a Fragment Mass Tolerance of 0.02 Da. Oxidation of methionine and carbamidomethylation of cysteine were specified as variable modifications for both search engines. All other program settings were left at default. Peptide spectral match (PSM) data, including associated PSM intensities, were exported from Proteome Discoverer, and the majority of further calculations and data processing were performed in Microsoft Excel and RStudio using custom scripts.

Peptide intensities were calculated by summing the intensities of all PSMs matching a given peptide. Peptide-to-protein inference was performed by distinguishing between exclusive (unique) and ambiguous (shared) peptides. Exclusive peptides, which map uniquely to a single protein, were used for primary protein quantification. Ambiguous peptides, mapping to two or more proteins, were handled separately to avoid inflating protein intensities and to prevent false-positive identifications. This separation is consistent with established proteomics workflows such as MaxQuant/Perseus ^51,52^ and ProteinProphet ^53^, which rely on unique peptides for reliable quantitative summarisation while retaining shared peptides for protein group inference. Similar strategies have been adopted in label-free and isobaric quantification pipelines ^54,55^ and in statistical frameworks such as MSstats ^56^. Hereafter, separate datasets were generated for exclusive and ambiguous peptides. Although the ambiguous peptide dataset is presented in Supplementary dataset1, all peptide and protein data, unless stated otherwise, refer to the exclusive peptide dataset. At this stage, splice variants for a particular locus were collapsed into a single protein entry per locus.

Protein intensity data comparisons were performed using an in-house Shiny app, Manchester Proteome Profiler, which uses the DEP package as the underlying engine ^57^. Protein intensities were normalised using the “quantile” normalisation option in DEP, which aligns the distributions of feature intensities across all samples. Missing values were imputed using the “man” option within DEP, which draws random numbers from a left-shifted, truncated normal distribution. Default settings were used for “man” imputation. Ratios (log2 fold changes) and p-values were then calculated for all proteins in each comparison. In our seedling interactome, we have 10 test baits and 7 control baits. All 17 baits were compared with all 7 control baits, excluding self-comparisons. Comparison data were exported from DEP, and further data analysis and plotting were performed in Excel and RStudio. Fold changes and p-values from these comparisons were then used to generate the volcano plots shown throughout this manuscript.

A protein was deemed to be “enriched” in a given comparison if the p-value was <0.05 and the log2 fold-enrichment was >1. This was the baseline level of enrichment, and we sometimes categorised enrichment as high (A1; p-value <0.001, log2 fold-enrichment >3.32), medium (A2; p-value <0.01, log2 fold-enrichment >2.32) and low (A3; p-value <0.05, log2 fold-enrichment >1). The corresponding scores for ambiguous peptide data were designated as B1, B2 and B3, respectively.

To balance the detection of the maximum number of interactions with confidence in detecting a particular interaction, we used various criteria, applied in the following order: detection of a protein with at least one peptide in a given bait protein; enrichment of interaction vs NoBait; enrichment of interaction vs EYFP; enrichment of interaction vs organelle reporter baits – BET12 (Golgi), STIM1 (ER), PIP2 (PM), MAP4 (microtubules) and LifeAct (actin). This was combined into a single score (1 to 5) based on the number of reporter baits against which an interaction is enriched.

All volcano plots in the manuscript include proteins that meet the first 3 criteria. Proteins are shown as colour-coded dots, based on enrichment relative to organelle reporter baits – green (enriched vs 5 organelle control baits), purple (enriched vs 4 organelle control baits), blue (enriched vs 3 organelle control baits), yellow (enriched vs 2 organelle control baits), brown (enriched vs 1 organelle control bait) and red (enriched vs 0 organelle control baits). *Functional enrichment analysis* Functional enrichment analysis was conducted in R using the clusterProfiler package ^58^ within the Bioconductor framework. The aim was to identify biological terms over-represented in a given protein list (preys for a particular bait). Input data comprised a spreadsheet containing protein identifiers, group (baits), and annotation terms (SUBA5 localisation). Terms were parsed and standardised using dplyr, tidyr, and stringr. A term-to-gene mapping (TERM2GENE) was constructed for enrichment analysis.

To avoid bias, enrichment was tested against a custom background universe comprising all genes detected in the experiment (or an optional global background file). Over-representation analysis (ORA) was performed using enricher() from clusterProfiler, which applies the hypergeometric test (equivalent to Fisher’s exact test) to assess whether the overlap between the group proteins and the annotation terms exceeds random expectation. P-values were adjusted for multiple testing using the Benjamini–Hochberg procedure to control the false discovery rate ^59^. Terms with adjusted p-values below a defined threshold (FDR < 0.05) were considered significantly enriched.

Analyses were conducted in R (version 4.5.2) using the packages clusterProfiler, ggplot2, dplyr, tidyr, readxl, and tidytext. All code and parameters are available upon request to ensure reproducibility.

### CESA antibody generation and testing

Peptide antibodies were synthesised and purified by the commercial provider LifeTein, LLC (New Jersey, USA). Peptides for CESA1 (C-SVRTTSGPLGPSDRNAISSPYID), CESA3 (C-KEKISERMLGWHLTRGKGE and C-GKRLPYSSDVNQSPNR), and CESA6 (C-NGIGFDQVSEGMSISRRN and C-RVHPVSLSDPTVAAHPR) were synthesised, and each peptide was injected into 2 rabbits. A total of 10 sera and antigen-affinity-purified pAb samples (5 peptides and 2 rabbits) were provided by the supplier. The primary antibodies were reconstituted to a concentration of 1 mg/mL. They were then tested and validated on Western blots using 6 crude protein extracts from 7-day-old seedlings of 6 different transgenic lines (2 independent lines each for *cesa1^any^*^1^, pCESA1::GFP-CESA1; *cesa3^ixr^*^1– 2^, pCESA3::GFP-CESA3; and *cesaC^prc^*^1–1^, pCESA6::GFP-CESA6). The untagged (endogenous) and GFP-tagged (transgenic) CESAs can be easily distinguished on the blots (Figure S16) by their distinct sizes. CESA1, CESA3, and CESA6 antibodies specifically detected the corresponding transgenic CESA band in the respective transgenic lines. The endogenous CESA band, on the other hand, was correctly detected in all samples. The only exception was the CESA6 antibodies, which detected the endogenous CESA in the CESA6 transgenic lines based on the *prc1-1* background. This indicates that our CESA6 antibodies can differentiate between CESA6 and CESA2/5/9. We also used a positive control, anti-GFP antibody, and a negative control (no primary antibody). The testing and validation results show that the antibodies are specific to the antigen-CESA and were used at a 1:1000 dilution.

### Protein expression analysis

7-day-old seedlings were harvested and placed in pre-weighed 1.5 ml screw-capped tubes containing a steel ball, then snap-frozen in liquid nitrogen. The tubes were re-weighed to determine tissue weight. The seedlings were then homogenised using a Qiagen TissueLyserII mixer mill with pre-frozen holding blocks to keep the tissue frozen. Laemmli buffer was added at 5 µl per mg of tissue powder. The samples were heated to 70°C for 20 minutes, then centrifuged at 14000 rpm in a benchtop centrifuge. The supernatant containing the crude protein extract was transferred to a new tube. The crude protein extracts were separated on homemade 7.5% polyacrylamide gels containing 0.1% trichloroacetic acid to facilitate stain-free imaging of the gels and membranes. The stain-free images were used as loading controls to normalise the signal from ECL images.

### Propidium-iodide staining

Arabidopsis etiolated hypocotyls from Col0 WT and the *bag1 bag2 bag3* triple mutant were stained by incubation in a 10 ug/mL solution of propidium iodide for 30 minutes. Seedlings were rinsed with water and mounted on glass slides. Confocal microscopy was performed on a Leica SP8 microscope using an HCX IRAPO L 25x/0.95 WATER objective. Samples were illuminated with a 588 nm laser, and light was collected in the 617–648 nm range. Images of complete hypocotyls were obtained by taking multiple overlapping images along the hypocotyls using the navigator mode in LASX software. Hypocotyls were straightened using the straighten function in Fiji, and the number of cells along a cell file was counted.

### qPCR Analysis

For transcript analysis, total RNA was extracted from whole 8-day-old Arabidopsis thaliana seedlings using the Bio-Rad Aurum Total RNA extraction kit, including on-column DNAase digestion. Following quality control, 1 ug of total RNA was converted into cDNA using the iScript cDNA synthesis kit, and the resulting cDNA was diluted 1:10 with nuclease-free water to serve as the template for subsequent amplification. The qPCR assays were performed in a 15 ul reaction volume containing 0.2 µM primers, 2 µl of cDNA template, and 7.5 ul of 2x qPCRBIO Lo-ROX SYBR green master mix (PCR Biosystems, Catalog# PB20.11-05). Primers (Table S3) were designed to span exon-exon junctions to preclude amplification of any residual genomic DNA. PCR amplifications were performed on a QuantStudio 5 qPCR machine (Applied Biosystems). Thermal cycling conditions included an initial denaturation at 95°C for 2 minutes, followed by 40 cycles of 95°C for 15s, 58°C for 30s, and 72°C for 15s. To verify primer specificity, a melting curve analysis was performed immediately after the amplification cycles, scanning from 65°C to 95°C. Relative transcript abundance was calculated using the 2^-DeltaDeltaCt^ method. Target gene expression was normalised against the mean expression of 2 reference genes, EF1alpha (AT5G60390) and UBC9 (AT4G27960). Each biological sample was represented by 4 independent biological replicates, with three technical replicates performed for each qPCR reaction to account for pipetting variance.

### Live cell imaging

For live-cell imaging of YFP-CESA6 in bag mutants, the YFP-CESA6 line (a kind gift from Staffan Persson) was crossed into *the bag1 bag3* double mutant. Live-cell imaging of etiolated hypocotyls was performed on an inverted Andor Dragonfly spinning disc system, following the protocol described previously ^60^. Briefly, 3-day-old etiolated hypocotyls were mounted in a “sandwich setup” comprising a custom-made metal slide with a central 18-mm hole, a 22 mm x 32 mm rectangular coverslip at the bottom, and a thin agarose pad (made by placing a drop of 0.1% agarose on a 16-mm round coverslip and placing another round coverslip on top). Imaging was performed using either a 60×oil-immersion objective (NA 1.4) or a 100× oil-immersion objective (NA 1.4). The EYFP fluorophore was excited with a 488 nm solid-state laser at 50% intensity, and emission was collected through a 525 filter. Time-lapse sequences were acquired as single-plane images using an Andor Sona 6 or Zyla sCMOS camera. Frames were captured at 10-second intervals over a total duration of 10 minutes (60 frames), with a 500 ms exposure time.

Quantification of CSC motility was performed using the standardised pipeline described by Verbančič et al. (2021). Raw time-lapse images were first processed in Fiji/ImageJ to subtract the background using a rolling ball radius of 50 pixels, correct for lateral drift using the StackReg plugin if needed, and apply a 4-frame walking average. To isolate the movement of plasma membrane-localised CSCs, sum-intensity projections were generated to reveal CSC tracks, which were then manually marked with segmented-line ROIs. CSC speeds were determined by generating kymographs along the trajectories of individual tracks. The slope of the resulting lines on the kymographs was measured to calculate the velocity of each CSC. Multiple tracks from multiple seedlings were analysed.

### Imaging of YFP-CESAC in ColO and bag13 mutant in light-grown roots

Live-cell imaging of root epidermal cells from 5-day-old light-grown seedlings was performed on a Zeiss LSM880 Airyscan system using a 40x (NA 1.2) water-immersion objective. YFP-BAG1 and YFP-CESA6 in WT and *bag1,bag3* backgrounds were analysed on this platform. EYFP was excited with a 514 nm laser (10% laser power), and emission was detected using the Airyscan detector in “super resolution” (SR) mode.

### Concanamycin treatment

To assess endomembrane trafficking and intracellular aggregation, seedlings were treated with the V-ATPase inhibitor concanamycin A (ConcA). A 1 µg/mL (approx. 1.15 µM) working solution of ConcA was prepared by diluting a 1 mg/mL stock (Cayman Chemical, USA; Item No. 11050) in sterile deionised water. An equivalent volume of acetonitrile (0.1% v/v final concentration) served as the mock solvent control. Five-day-old light-grown seedlings were transferred to 24-well plates containing 1.5 mL of either the ConcA or mock control solution. Seedlings were incubated in the respective solutions for 1.5 or 3 hours before imaging. Confocal microscopy of the primary root elongation zone was performed on a Zeiss LSM880 Airyscan microscope. To ensure accurate visual and quantitative comparisons, all imaging parameters (including laser power, detector gain, and pinhole size) were kept strictly constant across all genotype and treatment combinations throughout the experiment. At least three independent seedlings were imaged per condition.

### Dark treatment of light-grown seedling

5-day-old light-grown Arabidopsis seedlings were subjected to an extended dark treatment. Seedlings were incubated in complete darkness for 0 (untreated control), 2.45, or 6 hours before imaging. Confocal microscopy was performed on a Zeiss LSM880 Airyscan microscope. To comprehensively assess intracellular YFP-CESA6 distribution along the developmental gradient of the primary root, images were acquired at three distinct spatial locations: the root tip (Location 1), approximately 500 µm from the root tip (Location 2), and approximately 1000 µm from the root tip (Location 3). For each genotype and treatment time-point combination, 5 to 7 independent seedlings were imaged. To ensure accurate visual and quantitative comparisons of fluorescence intensities, all imaging acquisition parameters (including laser power, detector gain, and pinhole size) were kept constant throughout the experiment.

To quantify the intracellular distribution of YFP-CESA6, confocal micrographs were processed with a custom-written automated batch-processing macro in Fiji/ImageJ ^61^. To ensure high-fidelity detection of protein localisation, raw images were first subjected to background subtraction using a rolling ball algorithm (radius = 50 pixels) and a Gaussian blur (sigma = 1.0) to reduce high-frequency noise. Thresholding was performed using the IsoData algorithm to generate binary masks, followed by a watershed transformation to resolve closely apposed structures.

Particles were classified into four spatial categories based on their morphometric and densitometric properties. Mature Golgi bodies were identified by an area of 1.0–2.5 µm², consistent with established ultrastructural dimensions for plant Golgi cisternae ^62–64^. Aberrant protein accumulations were stratified by size into small (5.0–10.0 µm²), medium (10.0–15.0 µm²), and large (> 15.0 µm²) intracellular compartments.

To rigorously distinguish intracellular compartments from dense spatial clusters of wild-type Golgi stacks, a signal uniformity criterion was applied. The Coefficient of Variation (CV)—defined as the ratio of the standard deviation of signal intensity to the mean intensity—was calculated for each individual particle. Only structures with a CV < 0.3 were classified as true aggregates, as mutant-induced aggregations exhibit a highly homogeneous fluorescent signal, whereas clustered organelles show heterogeneous, multifocal intensity profiles. Morphological constraints (e.g., circularity and roundness) were minimised to ensure the unbiased inclusion of pleomorphic aggregate architectures. Data were exported as synchronised CSV files and processed in Microsoft Excel and R to produce the plots.

### Bimolecular Fluorescence Complementation (BiFC) Assays

To assess in planta protein-protein interactions, Bimolecular Fluorescence Complementation (BiFC) assays were performed using a custom-designed single-plasmid system to ensure comparable delivery of all transgene components. The full-length coding sequence of CESA1 was fused to the N-terminal fragment of YFP (nYFP), while the coding sequences of BAG1, BAG2, and BAG3 were individually fused to the C-terminal fragment of YFP (cYFP) using Gateway cloning. To circumvent variable co-transformation efficiencies, the nYFP and cYFP expression cassettes, together with a p19 viral silencing suppressor cassette to enhance transient expression, were assembled into a single pCambia-based binary vector backbone. Transcription of all three cassettes was driven by the CaMV 35S promoter.

The resulting sequence-verified single-plasmid BiFC constructs were transformed into *Agrobacterium tumefaciens* strain GV3101. Recombinant Agrobacterium cultures were harvested by centrifugation and resuspended in infiltration buffer (10 mM MgCl₂, 150 µM acetosyringone). As a positive control for infiltration and expression, the BiFC strains were mixed with an Agrobacterium strain harbouring a microtubule marker (pUBQ10::mCherry-MBD_MAP4). The suspensions were incubated at room temperature for 3 hours before co-infiltration into the abaxial epidermis of 3- to 4-week-old *Nicotiana benthamiana* leaves using a needleless syringe.

After infiltration, the plants were maintained under standard conditions for 72 hours. Leaf discs were excised from transiently expressing leaves and imaged on a Leica TCS SP8 upright confocal laser scanning microscope. The reconstituted BiFC (YFP) signal was excited with a 514 nm Argon laser, and emission was collected between 525 and 550 nm. The mCherry-MBD_MAP4 signal was excited with a 561 nm solid-state laser, and emission was collected between 580 and 620 nm. As a negative control, nYFP-CESA1 was paired with cYFP-AT5G18500. AT5G18500 is a membrane-localised protein in our broader proteomic dataset that does not interact with the CESAs. These negative control leaves were imaged under identical microscope settings.

## Author Contributions

M.K. and S.T. conceived the study. M.K. performed the experiments and analysed the data. S.T. contributed to experimental design and data interpretation and supervised the research. S.T. wrote the manuscript with input from M.K.

## Acknowledgements

We thank Xiaonan Lin, Wei Ching Tan and Shadab Farhadi for technical assistance. This work was supported by the Biotechnology and Biological Sciences Research Council (BBSRC) grant BB/X016919/1 awarded to S.T. Support M.K. was also provided through the Leverhulme Trust Research Project Grant RPG-2020-257 awarded to The University of Manchester.

