## supplementary figures and tables for "A cellulose synthase interactome uncovers BAG proteins as regulators of cellulose synthase homeostasis"

| Bait_type | Bait_Name | Bait_ID | Promoter | Bait_tag | Reference |
| --- | --- | --- | --- | --- | --- |
| Test | CESA1 | AT4G32410 | CESA1 | TurboID | (Arioli et al. 1998) |
| Test | CESA3 | AT5G05170 | CESA3 | TurboID | (Scheible et al. 2001) |
| Test | CESA6 | AT5G64740 | CESA6 | TurboID | (Fagard et al. 2000) |
| Test | KOR | AT5G49720 | KOR | TurboID | (Vain et al. 2014) |
| Test | CC1 | AT1G45688 | CC1 | TurboID | (Endler et al. 2015) |
| Test | CC2 | AT5G42860 | CC2 | TurboID | (Endler et al. 2015) |
| Test | CMU1 | AT4G10840 | CMU1 | TurboID | (Liu et al. 2016) |
| Test | CMU2 | AT3G27960 | CMU2 | TurboID | (Liu et al. 2016) |
| Test | SHOU4 | AT1G78880 | SHOU4 | TurboID | (Polko et al. 2018) |
| Test | SHOU4L | AT1G16860 | SHOU4 | TurboID | (Polko et al. 2018) |
| Control | EYFP | NA | CESA6 | TurboID.EYFP | NA |
| Control | STIM1 | NP_003147 | CESA6 | TurboID.EYFP | (Wu et al. 2006) |
| Control | BET12 | AT4G14455 | CESA6 | TurboID.EYFP | (Bolaños-Villegas et al. 2015) |
| Control | PIP2 | AT3G53420 | CESA6 | TurboID.EYFP | (Kammerloher et al. 1994) |
| Control | MAP4 | NP_002366.2 | CESA6 | TurboID.EYFP | (Marc et al. 1998) |
| Control | LifeAct | QHB11830.1 | CESA6 | TurboID.EYFP | (Riedl et al. 2008) |

Table S1. List of bait proteins used in this study

| Bait | Interactions detected | Interactions enriched vs NoBait | Interactions enriched vs NoBait+EYFP | Interactions enriched vs NoBait+EYFP+ 5/5 organelle baits |
| --- | --- | --- | --- | --- |
| **CESA1** | 597 | 11 | 49 | 16 |
| **CESA3** | 413 | 63 | 23 | 7 |
| **CESA6** | 547 | 122 | 66 | 27 |
| **KOR** | 992 | 339 | 214 | 63 |
| **CC1** | 1163 | 395 | 238 | 29 |
| **CC2** | 517 | 86 | 21 | 2 |
| **CMU1** | 1044 | 374 | 204 | 36 |
| **CMU2** | 632 | 105 | 47 | 9 |
| **SHOU4** | 1110 | 450 | 286 | 36 |
| **SHOU4L** | 893 | 188 | 87 | 2 |
| **NoBait** | 597 | 0 | 0 | 0 |
| **EYFP** | 1069 | 414 | 0 | 0 |
| **STIM1** | 1778 | 928 | 619 | 0 |
| **BET12** | 680 | 252 | 141 | 0 |
| **PIP2** | 1161 | 498 | 333 | 0 |
| **MAP4** | 968 | 330 | 133 | 0 |
| **LifeAct** | 1316 | 611 | 323 | 0 |
| **TOTAL Interactions** | **15477** | **5266** | **2784** | **227** |
| **Total Proteins** | **2748** | **1822** | **1333** | **152** |

**Table S2.** A summary table detailing the step-wise data filtering pipeline applied across all test baits. Starting from 2748 quantified proteins, the table tracks the number of interactions remaining after sequentially applying the base filtering criteria and subsequently filtering against varying stringency thresholds using the five organelle spatial reference controls (from enriched vs all 5/5 organelle baits down to 1/5). The resulting subsets of 2784 and 227 interactions (highlighted in red) were subsequently used to construct the interaction networks presented in Figure A2 and 2B, respectively.

| target | Forward primer sequence | Reverse primer sequence |
| --- | --- | --- |
| pCESA1 | gatGGCGGCCGgagaagagatggttaagaga | gatTCTAGA cgcagcaacgacaca |
| pCESA3 | gatGGCGGCCGctggatgatacagagtgatgg | gatTCTAGA ttgtcacttagttgtctc |
| pCESA6 | gatGGGCCCCgatgaatcaagttgaagtca | gatACTAGT atttctgtgaaacag |
| pKOR | gattGGTACC GGGGCCGgaagatggatggttcacaag | gatTCTAGA gatgatgctcttgata |
| pCC1 | gatGGCGGCCGcttcattggaaactctcac | gatTCTAGA tggtttcgatttgggaagtg |
| pCC2 | gatGGTACC GGGGCCGcttctgtctagcaaatccag | gatACTAGT ctggatgtggaatggaagat |
| pCMU1 | gatGGCGGCCGtaggttagaccagtggtgoc | gatACTAGT gaatgtgtctctctgtgggaag |
| pCMU2 | gatGGTACC GGGGCCGcaatatacaaaaataaaaatoc | gatACTAGT ggctccaaaactcacaactcaat |
| pSHOU4 | gatGGTACC GGGGCCGcttggttgttctcttttctc | gatTCTAGA ttgatttctcagggaactcgaactgg |
| pUBQ10 | gatGGGCCCC GGGGCCGcattcaaccatttatggata | gatACTAGT ctgttaatcagaaaaactcagtaaatc |
| pBAG1 | tgattacgaattcgagctcGGTACC CCGCAGG aattaaattttataaaatggta<br>gtcttgaag | ttgttacaacttgtACTAGT aatttttttagtaaaaaaactattgtctcttcc<br>ttgttacaacttgtACTAGT tttctttattaagagatagagagaaaaaacagag<br>gaatga |
| pBAG2 | tgattacgaattcgagctcGGTACC CCGCAGG cgtatcaatatagaacaaaag | ttgttacaacttgtACTAGT tcttgaatatttcttctctctctctctctct |
| pBAG3 | tgattacgaattcgagctcGGTACC CCGCAGG gcttctagtaagaagaat | ttgttacaacttgtACTAGT tcttgaatatttcttctctctctctctctctct |
| CESA1 | ggggacaaagmgtacaaaaaagcaggcttggATGGAAGCCAGTCCCG | ggggaccacmgtacaaagaagcctgggttCTAAAGACACCTCTCT |
| EYFP | ggggacaaagmgtacaaaaaagcaggcttggATGGTGAAGCAAGGGGAGG | ggggaccacmgtacaaagaagcctgggttTCACTTGTACAGCTGTCCA |
| TbID | ggggacaaagmgtacaaaaaagcaggcttggATGAAGACAATACTGTGCC | ggggaccacmgtacaaagaagcctgggttTACCTTTTGGCAGACCGCA |
| MAPPER | ggggacaaagmgtacaaaaaagcaggcttggATGGATGATGCGTCCGTCT | ggggaccacmgtacaaagaagcctgggttTCAAGTTACTGAATCTTCT |
| BET12 | caaaaaagcaggcttggATGAACCTTCGAAGGGAGAA | acaagaagcctgggttTACCCCTTGTATGTATTA |
| PIF2 | ggggacaaagmgtacaaaaaagcaggcttggATGGCAAGGATGTGAAGCC | ggggaccacmgtacaaagaagcctgggttTAGACGTTGGCAGCACTTCTGA |
| MAP4_MBD | ggggacaaagmgtacaaaaaagcaggcttggTCCCGCAAGAGAGCAAA | ggggaccacmgtacaaagaagcctgggttTCAACCTCTGCAGGAAAGT |
| LifeAct | ggggacaaagmgtacaaaaaagcaggcttggGCCACCATGGGCGTGGCCGAC<br>TTG | ggggaccacmgtacaaagaagcctgggttTCACTCTCTCTGGAGATGGACT<br>ACAAGAAAGCTGGGTATCAatcaagaatcccaat |
| BAG1 | caaaaaagcaggcttggATGATGAAGATGATGAGAA | ACAAGAAAGCTGGGTATTAagaataatcccaat |
| BAG2 | caaaaaagcaggcttggATGATGAAGATGATGATCGG | ACAAGAAAGCTGGGTATTAagaataatcccaat |
| BAG3 | caaaaaagcaggcttggATGATGAAGATGAATACAGG | ACAAGAAAGCTGGGTATTAatcgaagaatcccaat |
| Turbid-CESA1 | ctcagatctgctgagaagATGGAGGCCAGTGCCTGGC | GCCGGCACTGGCTCCATctctcagcagatctgag |
| Turbid-CESA3 | ctcagatctgctgagaagATGGAAATCCGAAGGAGAA | TTCCTCTGGGATCCATctctcagcagatctgag |
| Turbid-CESA6 | ctcagatctgctgagaagATGAACACCGGTGGTGGC | CCGACACCGGTGTTCTctctcagcagatctgag |
| Turbid-IRK2 | ctcagatctgctgagaagATGTACGGAAGATGCCA | TGGATCTCTCCGTACATctctcagcagatctgag |
| Turbid-CC1 | ctcagatctgctgagaagATGCACGCCAAACCCGAT | ATCGGTTTGGCGTGCATctctcagcagatctgag |
| Turbid-CC2 | ctcagatctgctgagaagATGCACGCCAAGACCCGAC | GTCGGTCTGGCGTGCATctctcagcagatctgag |
| Turbid-CMU1 | ctcagatctgctgagaagATGCCAGCAATGCCAGGT | ACCTGGCAITGCTGGCATctctcagcagatctgag |
| Turbid-CMU2 | ctcagatctgctgagaagATGGACGTAGGAGAGAGC | GCTCTCTCTAGTCCATctctcagcagatctgag |
| Turbid-SHOU4 | ctcagatctgctgagaagATGGGTTCCGAGATACCCA | TGGATCTCGAAGCCATctctcagcagatctgag |
| Turbid-SHOU4L | ctcagatctgctgagaagATGGGTTCCGAGATACCCA | TGGATCTCGAAGCCATctctcagcagatctgag |
| Turbid-EYFP | ctcagatctgctgagaagATGGTGAAGCAAGGGGAGG | CTCGCCCTTGGTCAACATctctcagcagatctgag |
| STIM1-Turbid | gacagcagcagcagcagcagATGAAGGATAATACTGTT | AACAGTATATCTTCActctcagcagatctgag |
| SM_3_15968 | TTTTCCGAGTATCGAGCTC | TTAGAATCAAGTACGGTGGCG |
| GABI_493F03 | CAACAATCACATTGCTGTCACCT | ACGTCTCCATCACTGAGATG |
| SALK_124153 | AGGTTGATTTTCCCATTC | CGAGTCTGTGCACTTCCGAAC |
| SALK_122722 | TGGTGGAGATTTTACCAG | CTTCTGTTCCGTAATCTCC |
| GABI_742B08 | ACCCGAAGGTACCCATAACTC | TCTCCGCAATTCGTTGAG |
| SALK_003242 | AGCCACCACTTAATCCCTTC | TCTACACGCGTACATCTAG |
| GABI_284G01 | GAGCACTGTGTCGATATCC | GAGTTCAAGCGGACATGAC |
| GABI_053H04 | TACCGAATGGAGTGTCCAG | TGAGTGGAGTGTATCTCTG |
| GABI_760D08 | GAGCTTTGTCAAGTCCCAATG | AACACACCCACGATTTCTTG |
| SALK_028810 | AATGACCGTTTACGCGGAC | CAAAAGCCCCAAATTTCTC |
| SALK_033390 | GATCAAGCTGATTTGACAGG | CTCAGCAACCTCACTGTCTCC |
| SALK_022140 | CTGCATAGGACCATGTCGAG | TATGCTTGGTAAATGACGGC |
| SALK_120883 | ATCCATGGCAATTTCTTAGG | ATCCATCAACCTCACTGCAAG |
| SAIL_764_H08 | GACCTTGAATCAAGGAAGCC | GTTTCTCTCTCCGACGATG |
| SALK_112105 | CATCAAGAAAGCGCAGAAC | CTGTTTGTCTTTGTTACGG |
| SALK_011984 | ACTTACCAATGCGTTCATGC | ATGTCCAGAGTTATAGCCGCC |
| SAIL_213_A09 | TGTCACATTTTGTCCCTTC | GACAGGACGATGAGGACTGC |
| SALK_020945 | CCTGTATCTGAGCCAGCAAG | CGTCTTGTACCAAGTTTTC |
| SAIL_829_G02 | ATTGTGTTGTTTGCACCTGG | CCAGGGCTCATAGAACACAG |
| SALK_046239 | ATGAAGCATCACTGGGAATG | ATCATCAGCAGCAGCCATATC |
| LB_SM | TACGAATAGAGCGTCCATTTAGAGTGA |  |
| LB_GABI-Kat | ATATTGACCATCACTCATTCG |  |
| LB_SalkB1.3 | ATTTTGGCGATTTCGGAAC |  |
| EF1alpha | TGTAAAGATGGATGCCACAC | GATCTGGTCAAGAGCCTCAAGAG |
| UBC9 | TCCTACTTCATGTAGCGCAGSAC | TCCTTCAGATGTGAGGCGAGATG |
| CESA1 | ATTGATCCAGCGCAACCTGTCC | TCTTCGCCATTGGAACCACTCC |
| CESA3 | ACCATCCAGGAATGATCCAGGTC | AACTTCTTCCGAGGTTTGGG |
| CESA6 | GTGCTCCCGAATGGATTTCTGC | CGTCACTTCCAGGAGACCTG |

**Table S3.** Primers used in this study
Restriction enzyme recognition sequences are shown in red, green and blue.

**A**

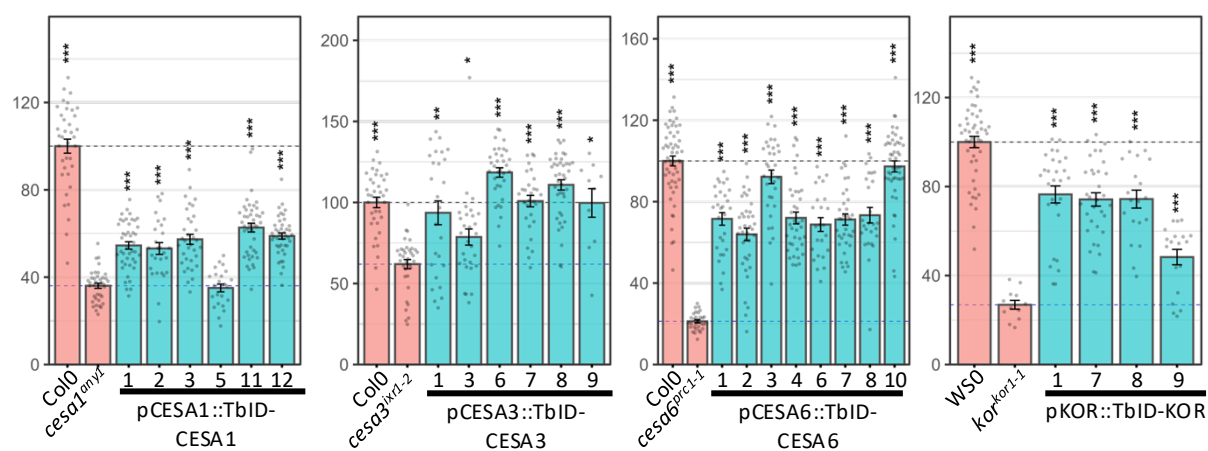

**B**

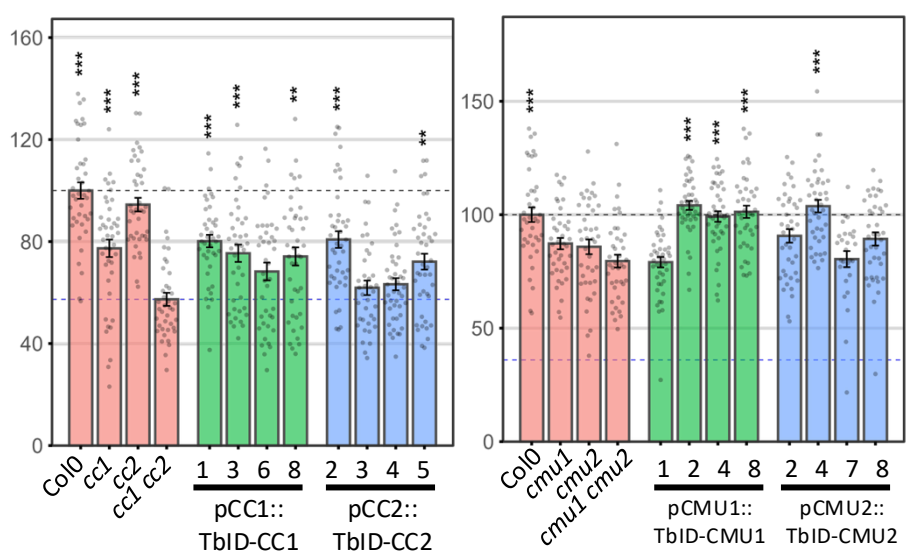

**C**

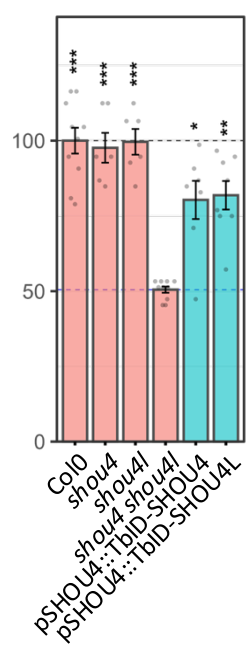

**Figure S1. Functional complementation analysis of TurboID-tagged test baits in corresponding mutant backgrounds.** To verify that the proximity labelling fusions retain their biological activity, TurboID (TbID)-tagged bait constructs were expressed in their respective Arabidopsis mutant backgrounds and assessed for phenotypic rescue. In all bar graphs, the black dashed line indicates the wild-type (WT) reference level (100%), and the red dashed line represents the mean measurement of the corresponding mutant background. Individual data points (open circles) represent biological replicates. **(A)** Quantification of etiolated hypocotyl length (expressed as % of WT) for seedlings grown on standard MS plates. The expression of *pCESA1::TbID-CESA1*, *pCESA3::TbID-CESA3*, *pCESA6::TbID-CESA6*, and *pKOR::TbID-KOR* successfully complements the hypocotyl elongation defects of the *cesa1*, *cesa3*, *cesa6*, and *kor* mutants, respectively. **(B)** Quantification of etiolated hypocotyl length (% of WT) for seedlings grown on plates supplemented with 300 nM oryzalin. The constructs *pCC1::TbID-CC1* and *pCC2::TbID-CC2* rescue the hypersensitivity of the *cc1 cc2* double mutant, while *pCMU1::TbID-CMU1* and *pCMU2::TbID-CMU2* complement the *cmu1 cmu2* double mutant. **(C)** Quantification of primary stem height (% of WT) in 7-week-old mature plants. The constructs *pSHOU4::TbID-SHOU4* and *pSHOU4::TbID-SHOU4L* effectively rescue the dwarfed phenotype of the *shou4 shou4l* double mutant.

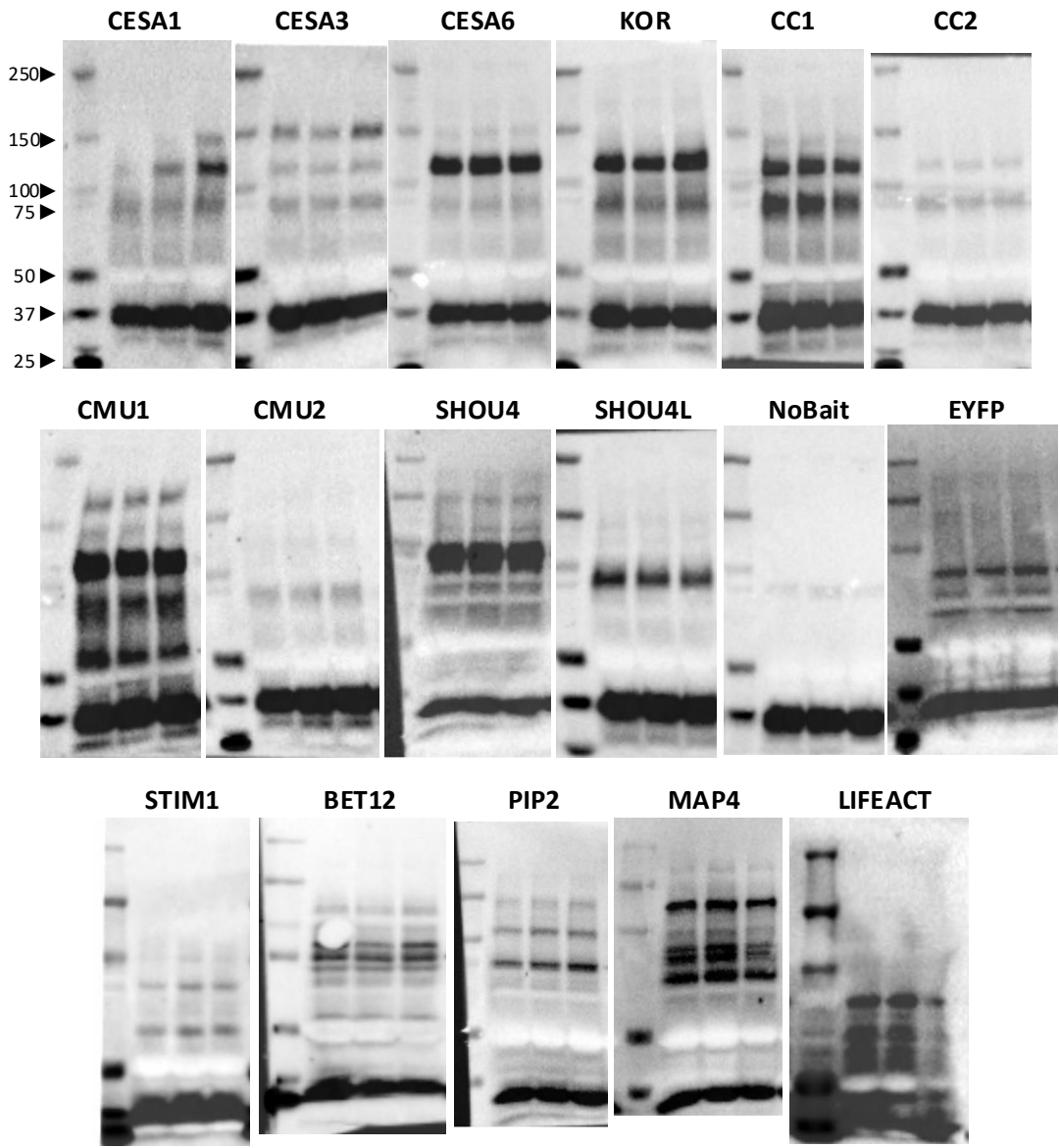

**Figure S2. Verification of proximity labelling reproducibility in biological triplicates.** To ensure the consistency of the samples prior to large-scale mass spectrometry analysis, biological triplicates of 8-day-old *Arabidopsis* seedlings expressing the indicated proximity labelling baits were treated with biotin for 3 h. Crude protein extracts from these three independent biological replicates were analysed by Western blotting using Streptavidin-HRP to detect the *in vivo* biotinylation patterns. Molecular weight markers (in kDa) are indicated on the left of the representative panel. Each panel displays the labelling profile of the triplicates for a specific test bait (e.g., CESA1, CESA3, KOR, CC1) or control line (e.g., NoBait, EYFP). The uniform banding patterns and signal intensities across the three lanes for each construct demonstrate highly reproducible proximity labelling suitable for quantitative multi-bait proteomics.

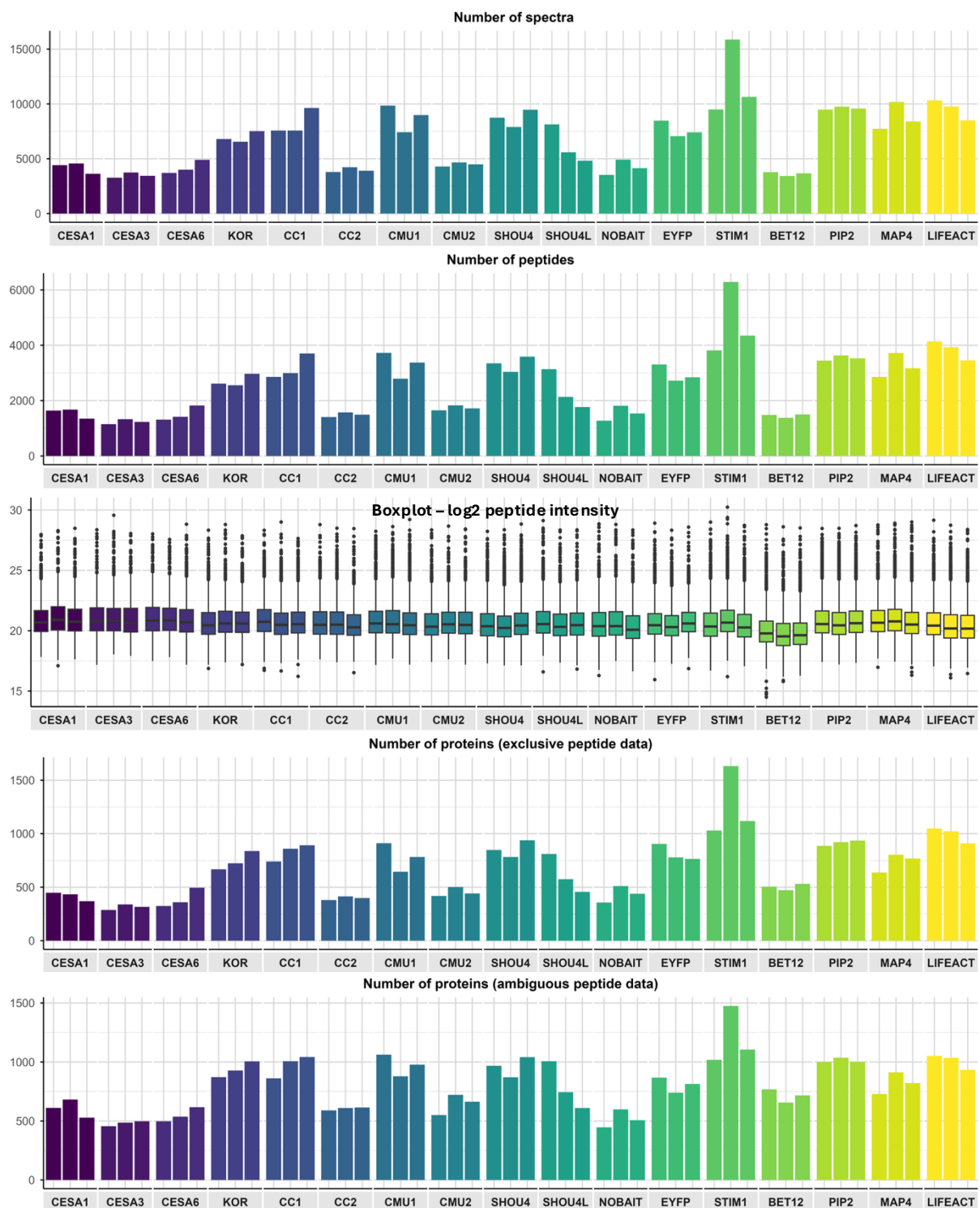

**Figure S3. Global overview and quality control of the multi-bait proximity labelling proteomics dataset.** The panels summarise the overall data yield and quantitative distribution across all biological triplicates for each test and control bait following LC-MS/MS analysis. From top to bottom, the charts display: the total number of peptide spectrum matches (PSMs, denoted as 'spectra'); the total number of identified peptides; a boxplot showing the distribution of log<sub>2</sub>-transformed peptide intensities across samples; the total number of identified proteins based strictly on unique peptides ('exclusive peptide data'); and the total number of identified proteins including those mapped with shared peptides ('ambiguous peptide data'). The consistent feature counts and intensity distributions across the three independent biological replicates for each bait demonstrate the robust reproducibility of both the *in vivo* proximity labelling and the mass spectrometry acquisition.

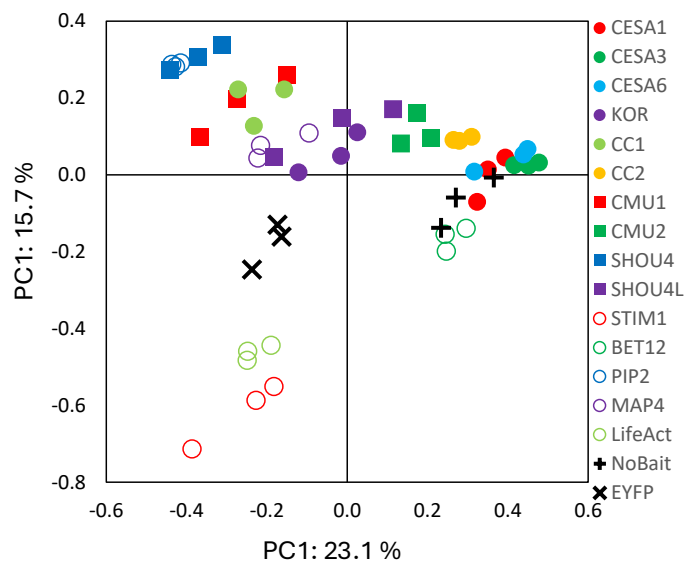

**Figure S4. Principal Component Analysis (PCA) of the proximity labelling mass spectrometry datasets.** A PCA scatter plot visualizing the variance and clustering of all 51 independent samples based on exclusive peptide data. The dataset encompasses three independent biological replicates for each of the 17 experimental conditions: the 10 Cellulose Synthase Complex (CSC) test baits (solid circles and squares), the 5 organelle spatial controls (open circles), and the 2 baseline negative controls (NoBait and EYFP; crosses and plus symbols). The tight clustering of replicates of the same color/shape demonstrates the high biological reproducibility of the mass spectrometry data. Furthermore, the distinct spatial separation between the CSC test baits and the control groups underscores the specificity of the proximity labelling approach and validates the distinct proteomic microenvironments captured by each bait category.

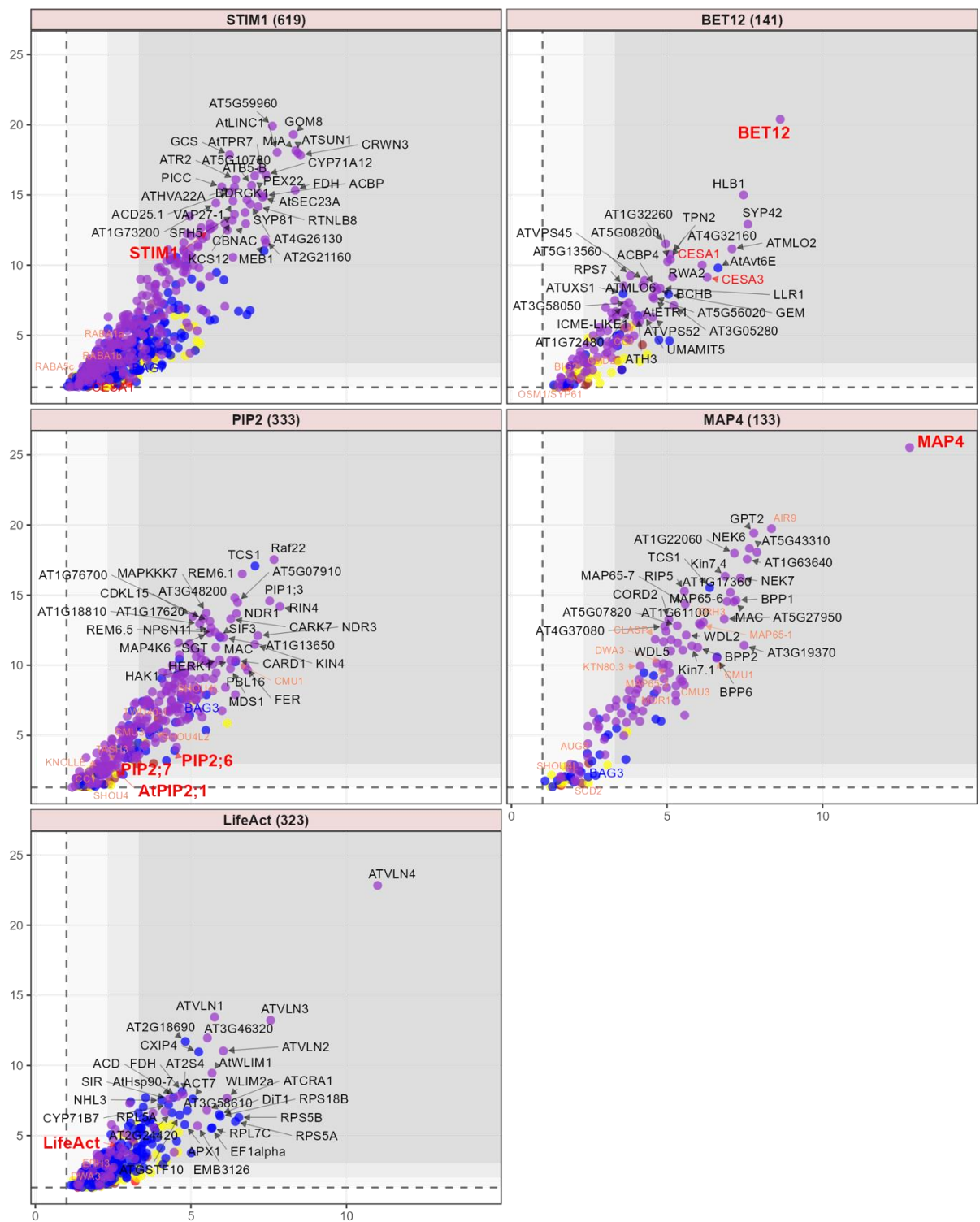

**Figure S5. Differential enrichment volcano plots for individual baits (exclusive peptide data).** Multi-panel volcano plots illustrating the differential protein enrichment of 1 test (blue panel headers) and 5 control (pink panel header) bait proteins versus the EYFP control. The total number of interactors for each bait is provided in parentheses within the panel headers. For each panel, proteins are plotted by their log<sub>2</sub> fold change on the x-axis and -log<sub>10</sub> *p*-value on the y-axis.

Background shading is used to delineate statistical and fold-change thresholds: no shading ($|\log_2\text{FC}| \geq 1.0$, $p \leq 0.05$), light grey ($|\log_2\text{FC}| \geq 2.32$, $p \leq 0.01$), and dark grey ($|\log_2\text{FC}| \geq 3.32$, $p \leq 0.001$). Data points are colour-coded based on the number of organelle marker baits (out of 5) against which they remained significantly enriched: 0 (red), 1 (brown), 2 (yellow), 3 (blue), 4 (purple), and 5 (green).

Key interactors are annotated with text labels according to the following scheme: the bait is highlighted in **large red text**; CESA proteins are in red; BAG family proteins are in blue; and known cellulose-synthesis-related proteins (as defined in Gu and Rasmussen, 2022) are highlighted in salmon.

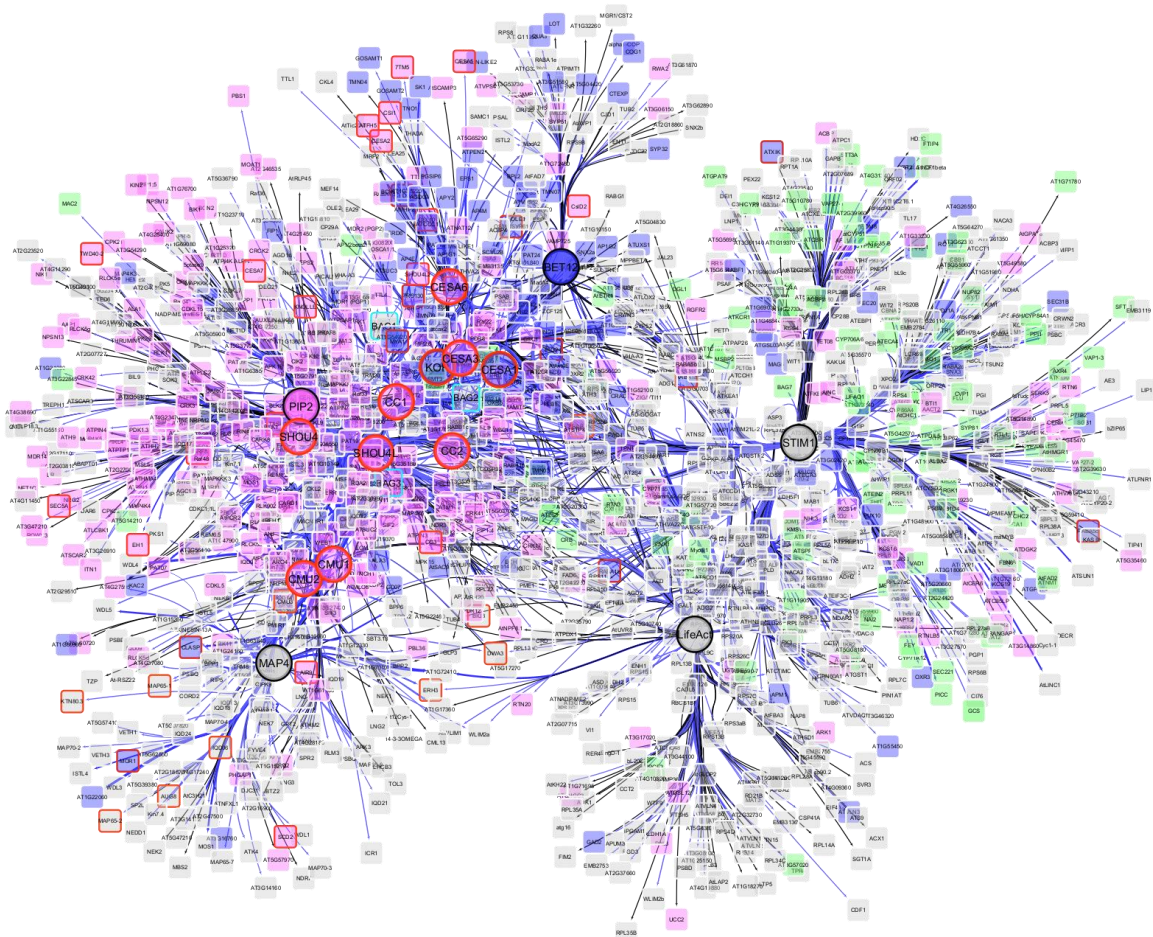

**Figure S6. The extended Cellulose Synthase Complex (CSC) proximity interaction network.** A comprehensive network graph visualising the 2,784 high-confidence proximity interactions identified following the base level of statistical filtering (significantly enriched against both NoBait and EYFP controls). Nodes represent individual identified proteins, and the connecting edges represent a significant proximity interaction with a specific bait. The large circular hubs represent the bait proteins used in the assay. Node core colours indicate subcellular localisation as annotated in the SUBA5 database (green: Endoplasmic Reticulum; magenta: Plasma Membrane; blue: Golgi apparatus). Nodes highlighted with a red border represent known cellulose-synthesis-related proteins (Gu and Rasmussen, 2022). Edge thickness is proportional to the number of unique peptides detecting the interaction. Edge colour also reflects this confidence metric: black edges indicate an interaction detected by a single peptide, while blue edges indicate an interaction detected by two or more unique peptides. The network was generated and visualised using Cytoscape, with edge bundling applied to enhance the clarity of complex interaction patterns.

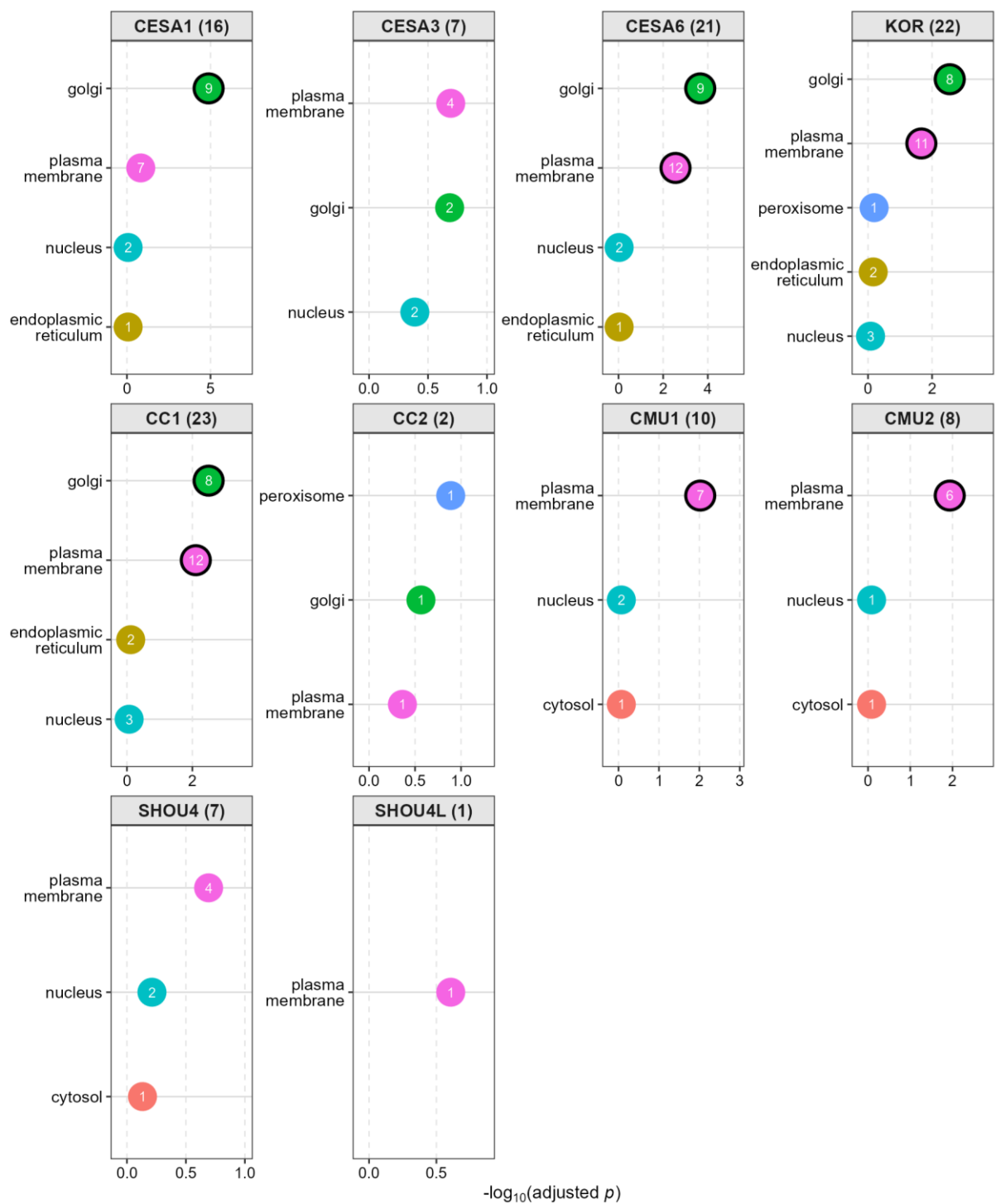

**Figure S7. Subcellular localization enrichment of the high confidence core interactome.** Scatter plots illustrating the top 5 most enriched subcellular localization categories (annotated via the SUBA5 database) for the 119 interaction network (Figure 3A). The x-axis indicates the statistical significance of the enrichment, expressed as the $-\log_{10}(\text{adjusted } p\text{-value})$. Each data point represents a specific subcellular compartment, with the number inside the circle indicating the gene count (the number of identified interactors mapping to that specific compartment). Data points featuring a solid black border denote statistically significant enrichment (adjusted $p < 0.05$), whereas points without a border indicate terms that did not reach statistical significance. The total number of interacting proteins analysed for each bait is indicated in parentheses within the individual panel headers.

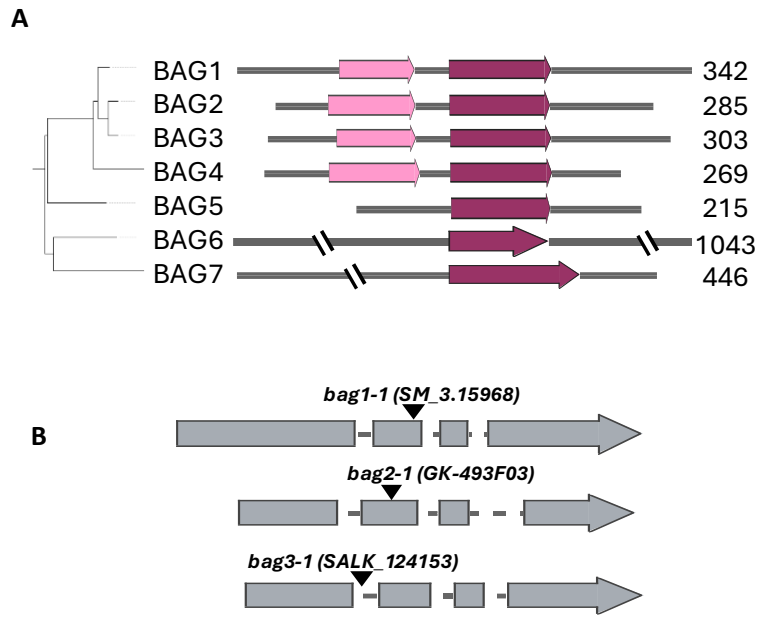

**Figure S8. Phylogeny, domain architecture, and mutant alleles of the *Arabidopsis* BAG protein family. (A)** Phylogenetic relationship and structural organisation of the seven *Arabidopsis thaliana* BAG proteins (BAG1–BAG7). The phylogenetic tree was generated from full-length protein sequences using a COBALT alignment and FastTree (midpoint rooted). Alongside the tree, schematic diagrams illustrate the conserved protein domains: the dark maroon regions represent the defining BAG domain present in all family members, while the light pink regions denote the N-terminal ubiquitin-like (UBL) domain, which is specific to the BAG1–BAG4 subclade. Total protein lengths in amino acids are indicated on the right. **(B)** Schematic representation of the *BAG1*, *BAG2*, and *BAG3* gene models showing the respective T-DNA insertion mutant alleles used in this study. Exons are depicted as grey boxes and introns as connecting lines. Black inverted triangles indicate the precise locations of the T-DNA insertions for the *bag1-1* (SM\_3.15968), *bag2-1* (GK-493F03), and *bag3-1* (SALK\_124153) knockout lines.

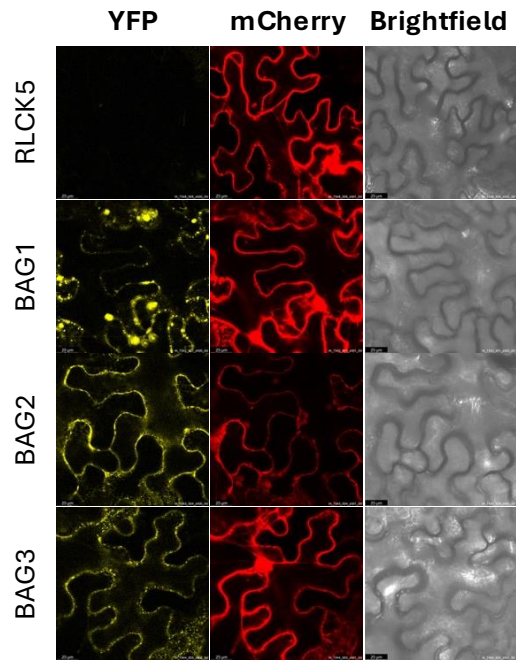

**Figure 9. Bimolecular Fluorescence Complementation (BiFC) assays demonstrating *in planta* interactions between BAG proteins and CESA1.** Confocal microscopy images of *Nicotiana benthamiana* (tobacco) leaf epidermal cells following transient co-expression of CESA1 fused to one half of the YFP fluorophore alongside BAG1, BAG2, or BAG3 fused to the complementary half. The reconstitution of the YFP signal indicates interaction between CESA1 and the BAG proteins *in vivo*. As a negative control for interaction specificity, CESA1 was co-expressed with RLCK5, an unrelated plasma membrane-localized protein not expected to interact with the CESA1, which resulted in no detectable YFP fluorescence. To confirm successful transformation and protein expression across all experiments, a plasmid containing mCherry-MAP4 was co-infiltrated as an internal control. Brightfield images were also captured to demonstrate the focal plane being imaged. A 25 micron scalebar is indicated in black.

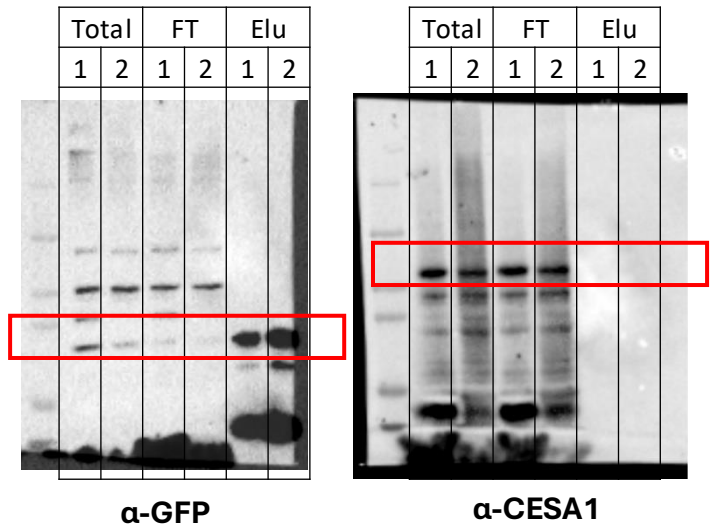

**Figure S10. Co-immunoprecipitation (Co-IP) analysis of HA-EYFP-BAG1 and endogenous CESA1.** Protein extracts from transgenic Arabidopsis seedlings expressing HA-EYFP-BAG1 (in a *bag1* mutant background) were subjected to immunoprecipitation using GFP-Trap magnetic beads. To optimise the solubilisation of the membrane-integrated Cellulose Synthase Complex (CSC), two different detergent lysis buffers were evaluated: Buffer 1 (1% DDM) and Buffer 2 (1% Triton X-100). Total protein extract (Total), unbound flow-through (FT), and bound elution (Elu) fractions were analysed by Western blotting. The left panel, probed with an -GFP antibody, confirms the successful immunoprecipitation and strong enrichment of the HA-EYFP-BAG1 bait protein (expected MW ~67.7 kDa, red box) in the elution fractions under both detergent conditions. The right panel, probed with CESA1 antibody, demonstrates robust detection of endogenous CESA1 in the Total and FT lanes. However, CESA1 failed to co-purify with BAG1 in the elution fractions. The inability to capture this complex using standard detergent-based Co-IP methods highlights the transient or structurally sensitive nature of the BAG1-CSC interaction during cell wall synthesis, thereby necessitating the use of *in vivo* proximity labelling approaches to capture this functional interactome.

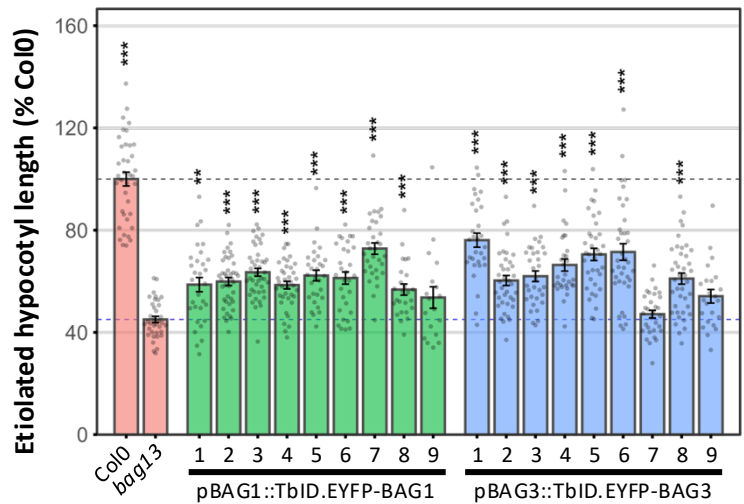

**Figure S11. Functional complementation of the *bag13* double mutant by TurboID-tagged BAG constructs.** Quantification of etiolated hypocotyl elongation in 5-day-old dark-grown seedlings cultivated on media containing 1 nM isoxaben. The *bag1 bag3* (*bag13*) double mutant exhibits hypersensitivity to this cellulose biosynthesis inhibitor, resulting in severely restricted hypocotyl elongation compared to the wild-type (Col0) control. To verify the biological functionality of the bait constructs utilised for proximity labelling, the *bag13* mutant was transformed with either *pBAG1::TbID-EYFP-BAG1* (green bars) or *pBAG3::TbID-EYFP-BAG3* (blue bars). Analysis of 9 independent transgenic lines for each construct demonstrates a significant restoration of hypocotyl elongation. This successful complementation confirms that the TurboID-tagged BAG fusion proteins retain their wild-type physiological function *in vivo*. Data are expressed as a percentage of the Col0 wild-type average length. Bar heights represent the mean, translucent dots represent individual biological replicates, and error bars denote the standard error of the mean (SEM). Asterisks indicate statistical significance compared to the uncomplemented *bag13* mutant background (\*\*\*) $p < 0.001$.

### Inhibitor Sensitivity - IXB-1nM

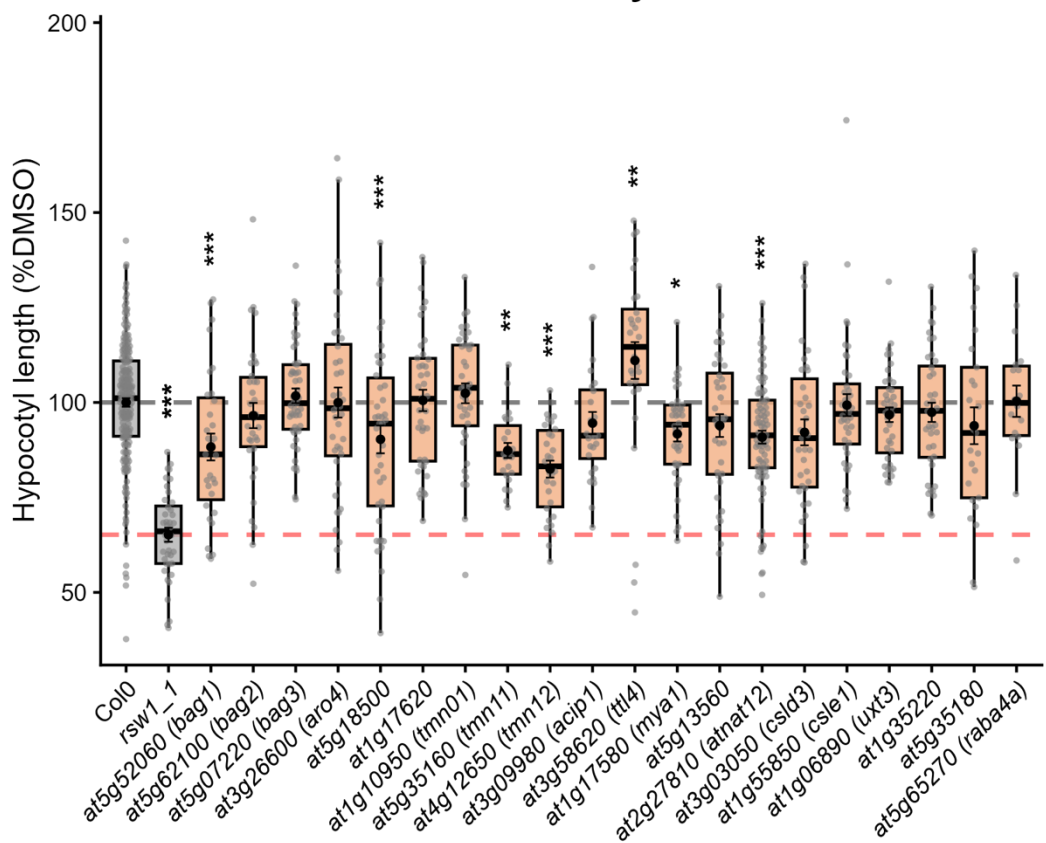

**Figure S12. Differential sensitivity of candidate interactome mutants to cellulose biosynthesis inhibitor isoxaben**

Seedlings were grown on plates containing 1 nM Isoxaben (IXB) or DMSO (mock) and etiolated hypocotyl lengths were measured. To isolate true chemical sensitivity from inherent genetic growth defects, data are double-normalized and expressed as a percentage of each specific genotype's own mean hypocotyl length on mock (DMSO) media.

The boxplots display the median and interquartile ranges, with individual biological replicates overlaid as solid grey dots. Black dots and solid vertical error bars denote the mean ± standard error of the mean (SEM). Horizontal dashed lines provide visual reference for the mean values of the Col0 wild-type baseline (black) and the *rsw1-1* positive control (red). Statistical significance was determined using a one-way ANOVA followed by Dunnett's post-hoc test against the Col0 reference group (\* $p \leq 0.05$, \*\* $p \leq 0.01$, \*\*\* $p \leq 0.001$).

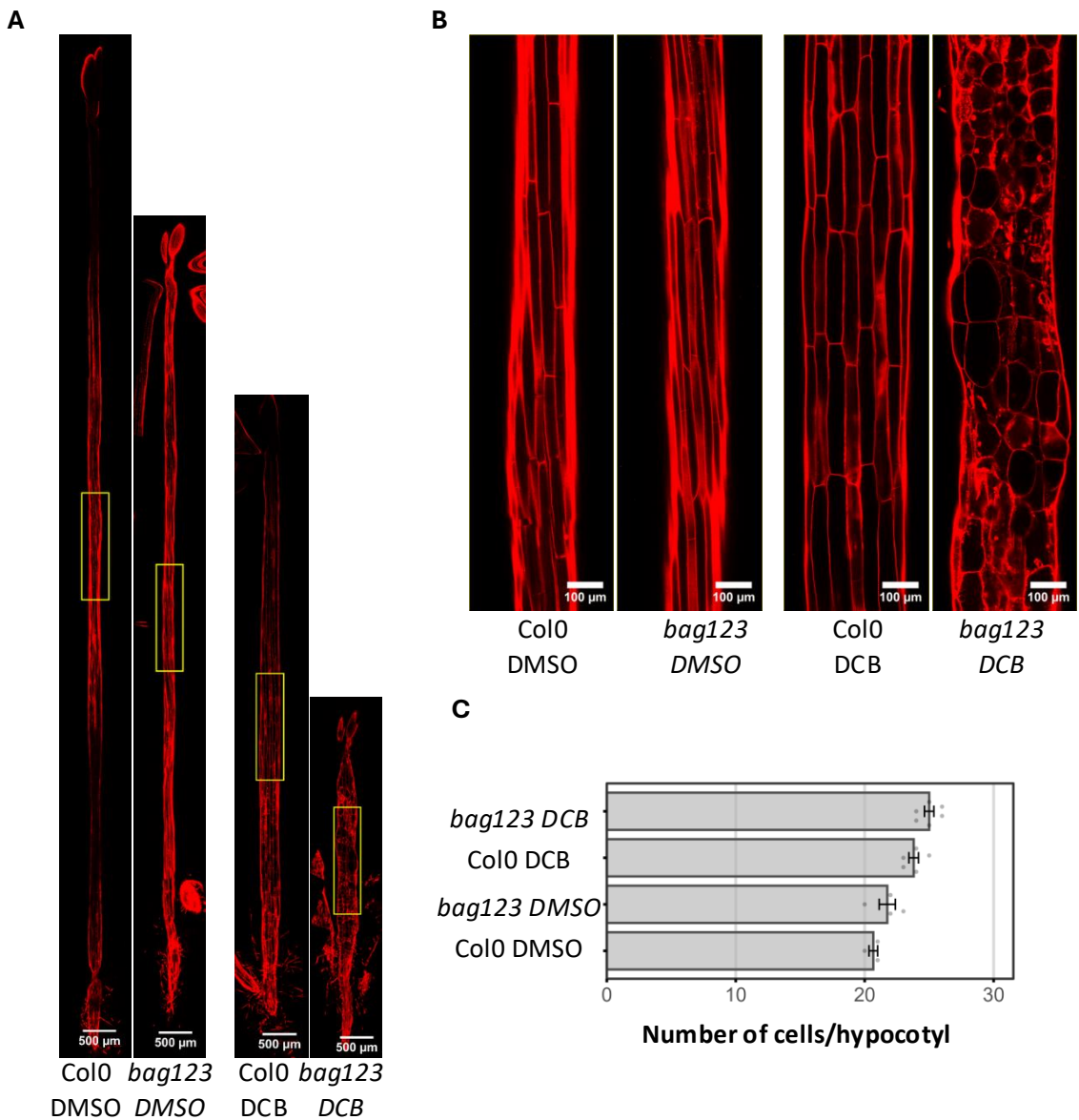

**Figure S13. Cellular dimensions of *Arabidopsis bag* etiolated hypocotyls.** (A) Representative full-length confocal images of 3-day-old etiolated hypocotyls grown on either DMSO control plates or 100 nM DCB plates. Seedlings of wild-type (Col0) and the *bag1/2/3* triple mutant were stained with propidium iodide to visualise cell walls and imaged using a Leica SP8 confocal microscope (561 nm laser excitation). Yellow rectangular boxes indicate the specific regions magnified in the subsequent panel. (B) High-resolution zoomed-in views of the selected areas from (A), detailing individual epidermal cell morphology and dimensions. (C) Quantification of the total number of cells counted along a single longitudinal epidermal file across the entire length of the hypocotyl for the indicated genotypes.

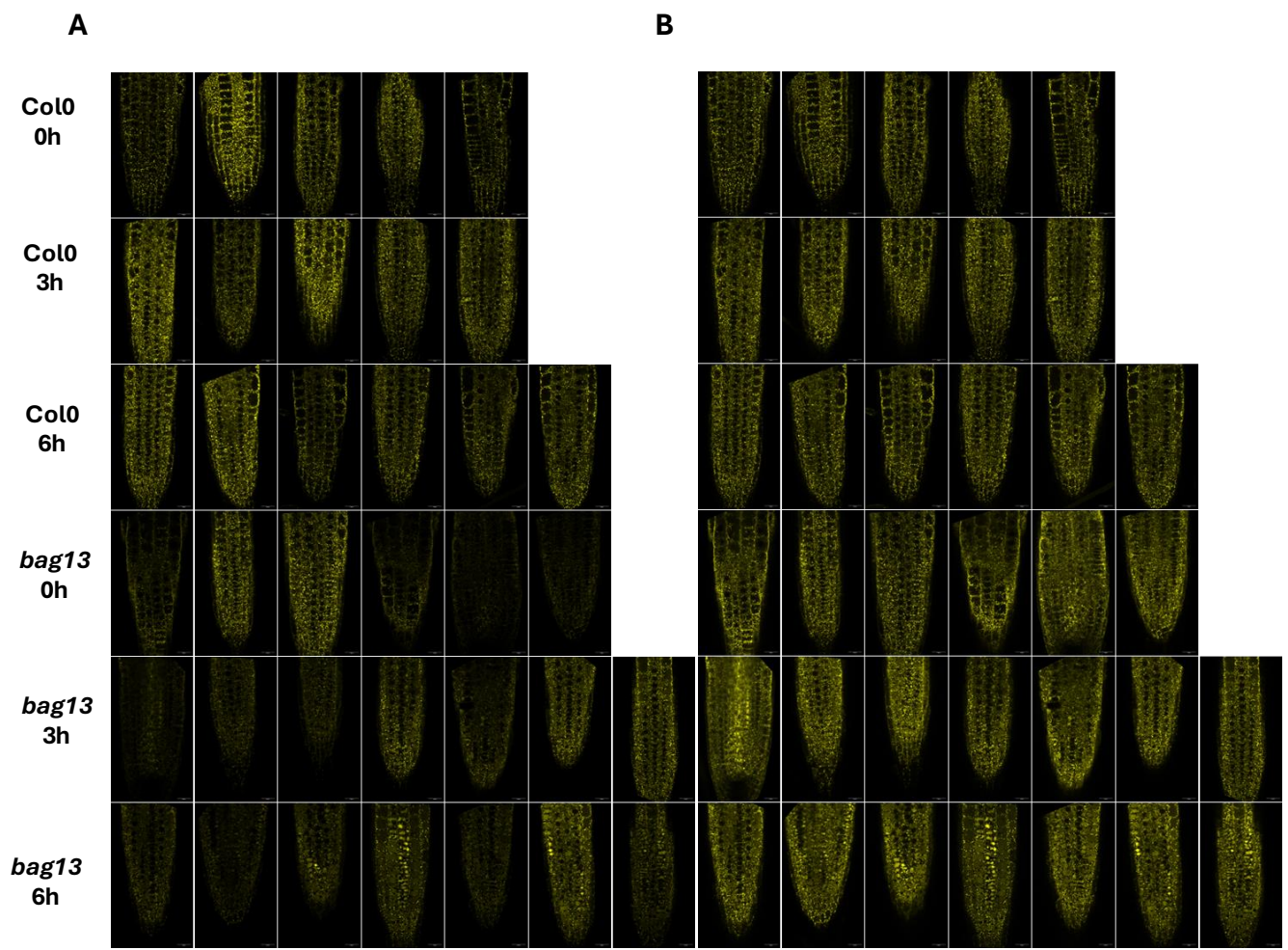

**Figure S14. Comprehensive spatial overview of dark-induced YFP-CESA6 internalisation.**

Confocal microscopy images of 5-day-old wild-type (Col0) and *bag1 bag3* double mutant seedlings expressing YFP-CESA6. Seedlings were incubated in complete darkness for 0, 2.45, or 6 hours. **(A)** Images displayed with strictly fixed brightness and contrast parameters (100–8000 display range) applied uniformly across all panels. This absolute scaling allows for direct visual comparison of overall signal intensities, revealing the differences in CSC abundance between the wild-type and the *bag1 bag3* mutant over the time course. **(B)** The identical image dataset as in (A), but with brightness and contrast individually optimised ('Auto' adjustment in Fiji/ImageJ) for each panel. This relative scaling maximises the dynamic range within individual images to clearly highlight spatial distribution and structural morphology, specifically emphasizing the pronounced vacuolar accumulation and intracellular aggregation of YFP-CESA6 in the *bag1 bag3* mutant following prolonged dark incubation.

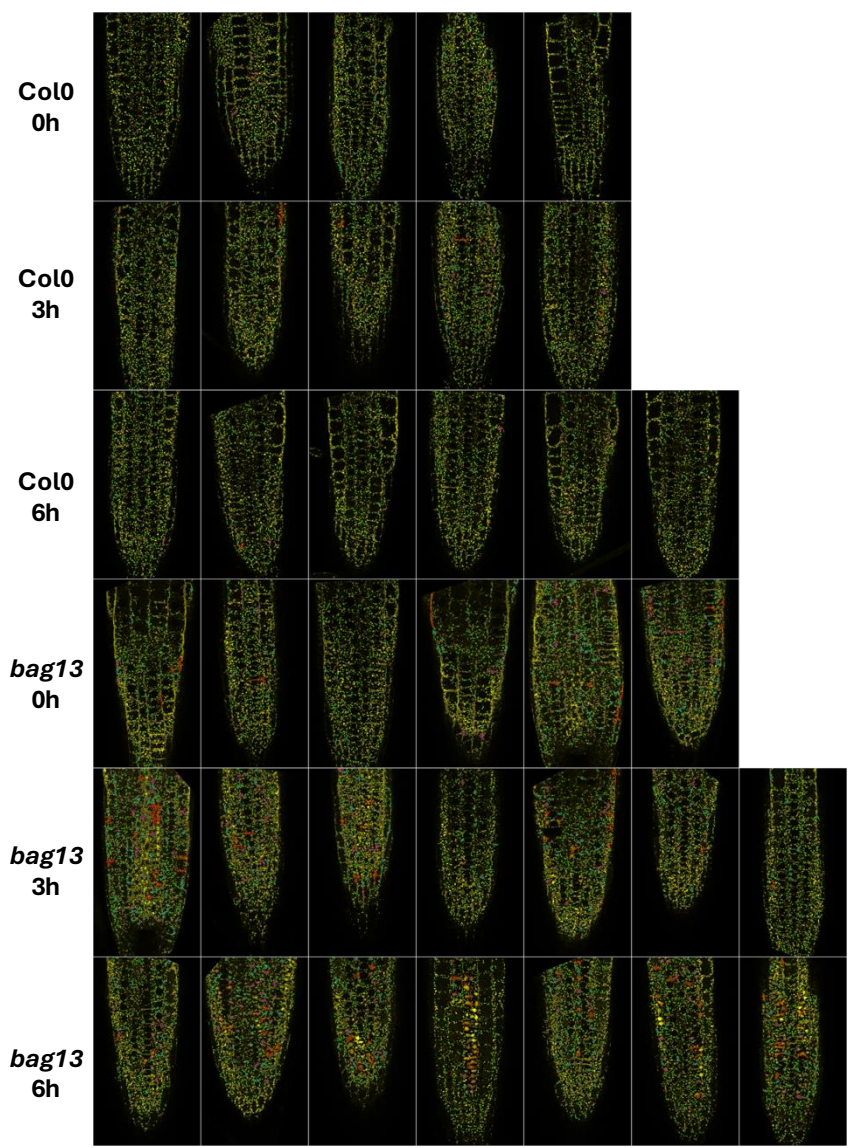

**Figure S15. Spatially resolved computational segmentation of dark-induced YFP-CESA6 internalisation.** Visual representation of the quantitative image analysis pipelines applied to 5-day-old wild-type (Col0) and *bag13* double mutant root epidermal cells following 0, 3, or 6 hours of dark treatment. For segmentation analysis, confocal images were subjected to rolling-ball background subtraction and IsoData auto-thresholding. Intracellular fluorescent signals were computationally segmented and classified based on particle area, solidity, and coefficient of variation (CV) in the signal intensity. The images display the resulting overlays, where particles are demarcated by uniquely coloured outlines corresponding to discrete structural classes: Large Aggregates (red outlines; Area > 15 $\mu\text{m}^2$, CV 0.3), Medium Aggregates (magenta outlines; Area 10–15 $\mu\text{m}^2$, CV 0.3), Small Aggregates (cyan outlines; Area 5–10 $\mu\text{m}^2$, CV 0.3), or Golgi-associated structures (green outlines; Area 1.0–2.5 $\mu\text{m}^2$). Quantification of large+medium aggregates is presented in Figure 6C.

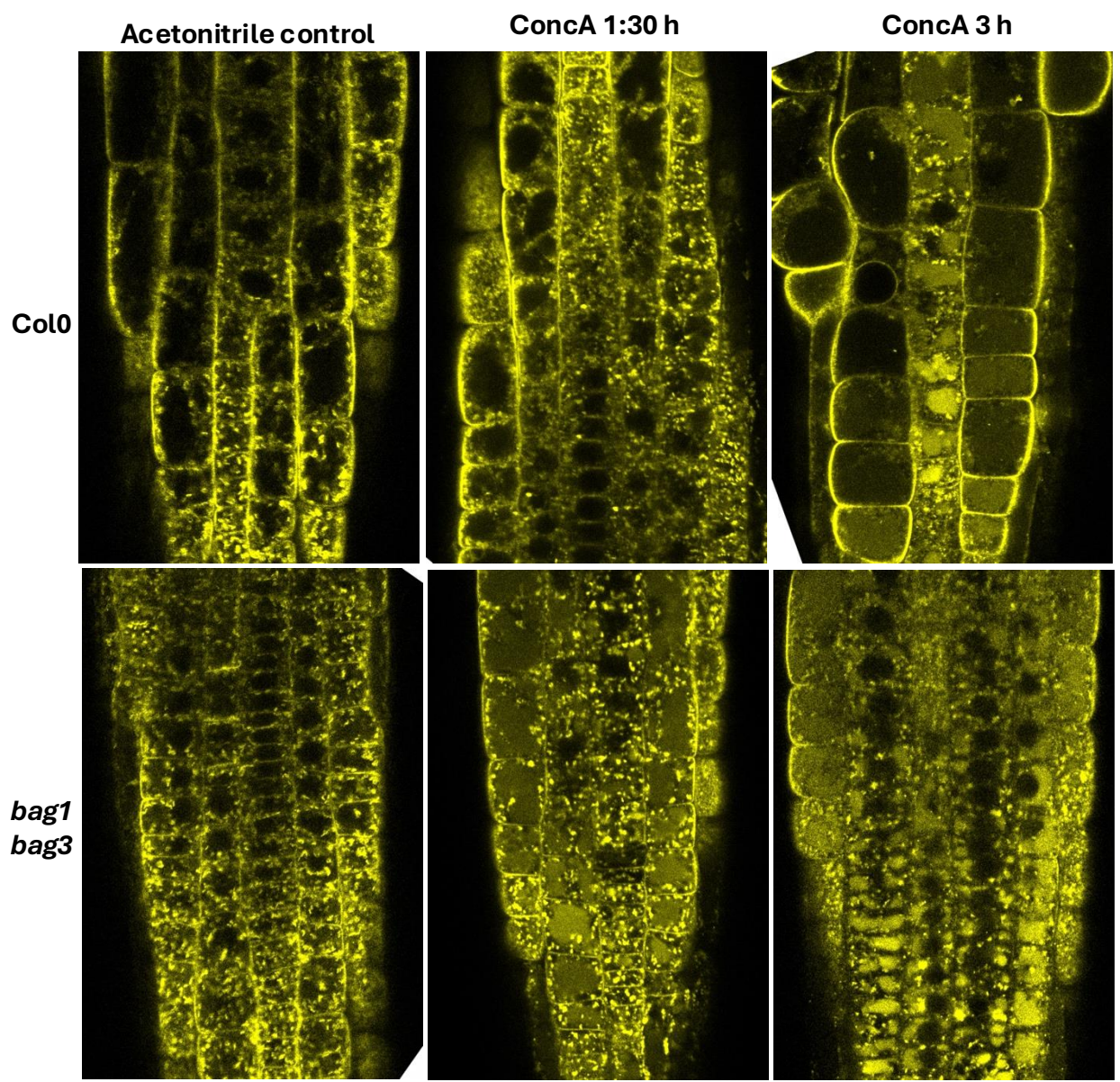

**Figure S16. Effect of the V-ATPase inhibitor concanamycin A on CSC intracellular trafficking in *bag* mutants.** Representative confocal images of root tip epidermal cells from 5-day-old light-grown wild-type (Col0) and *bag1 bag3* double mutant seedlings expressing YFP-CESA6 are shown. Seedlings were incubated in a liquid medium containing either an acetonitrile solvent control or 1 $\mu\text{g/mL}$ concanamycin A (ConcA) for 1.5 and 3 hours prior to imaging on a Zeiss Airyscan microscope. Imaging settings were kept strictly constant across all time points and genotypes to allow for direct visual comparison.

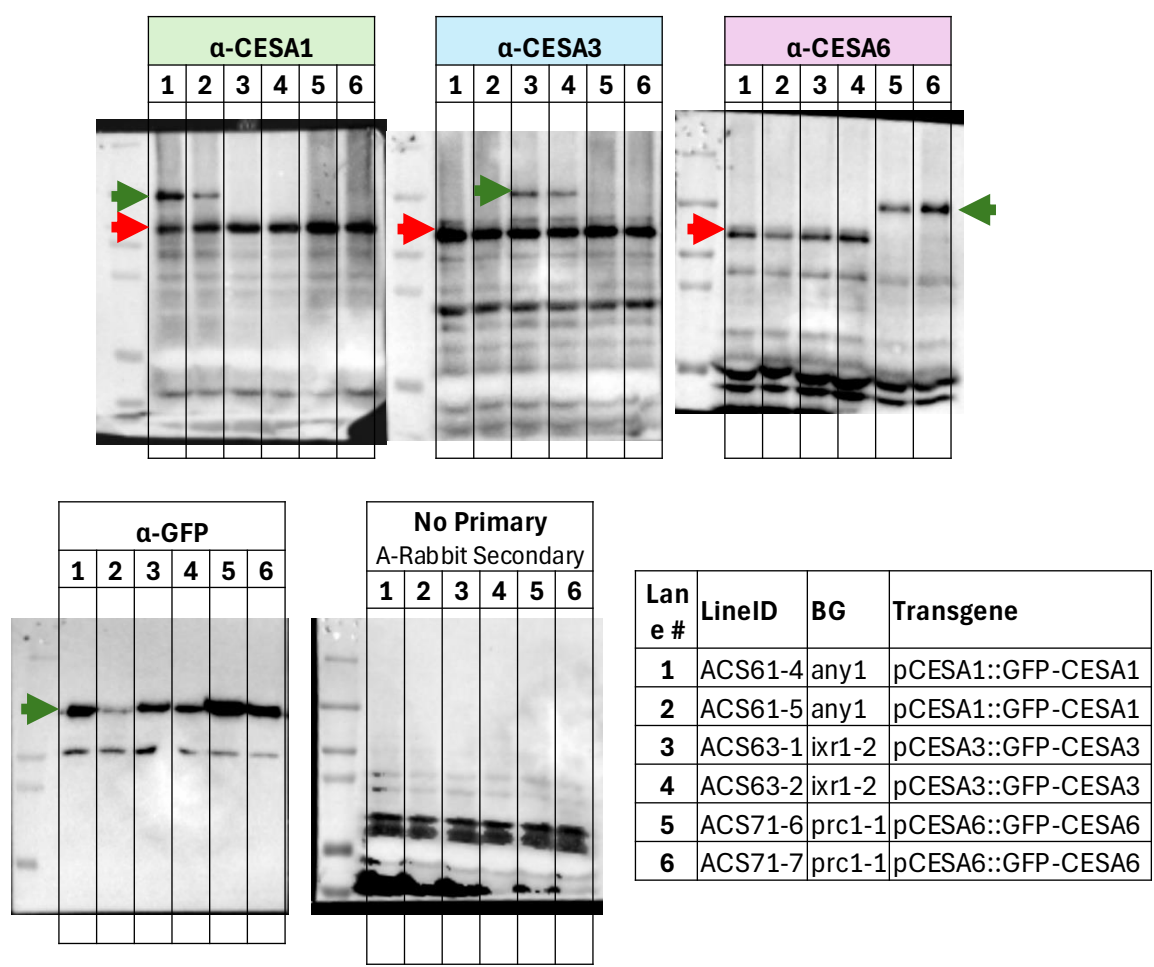

**Figure S17. Validation of custom isoform-specific Cellulose Synthase (CESA) antibodies.** Western blot analysis evaluating the specificity of custom rabbit polyclonal antisera raised against CESA1, CESA3, and CESA6. Total protein extracts were generated from two independent stable transgenic *Arabidopsis* lines expressing GFP-tagged CESA proteins in their corresponding mutant backgrounds: *pCESA1::GFP-CESA1* in *any1* (Lanes 1 and 2), *pCESA3::GFP-CESA3* in *ixr1-2* (Lanes 3 and 4), and *pCESA6::GFP-CESA6* in *prc1-1* (Lanes 5 and 6). Across the top panels, green arrows indicate the specific detection of the heavier, GFP-tagged CESA transgene. Red arrows denote the detection of the endogenous CESA protein. The complete absence of the endogenous CESA6 band (red arrow) in lanes 5 and 6 confirms the specificity of the α-CESA6 antibody, as these lines are in the *prc1-1* null mutant background. The bottom panels serve as procedural controls, comprising an immunoblot probed with an α-GFP antibody to independently verify the migration of the transgene (left), and a secondary-antibody-only blot to assess non-specific background binding (right).
